# Comprehensive epitope mutational scan database enables accurate T cell receptor cross-reactivity prediction

**DOI:** 10.1101/2024.01.22.576714

**Authors:** Amitava Banerjee, David J Pattinson, Cornelia L. Wincek, Paul Bunk, Armend Axhemi, Sarah R. Chapin, Saket Navlakha, Hannah V. Meyer

## Abstract

Predicting T cell receptor (TCR) activation is challenging due to the lack of both unbiased benchmarking datasets and computational methods that are sensitive to small mutations to a peptide. To address these challenges, we curated a comprehensive database, called BATCAVE, encompassing complete single amino acid mutational assays of more than 22,000 TCR-peptide pairs, centered around 25 immunogenic human and mouse epitopes, across both major histocompatibility complex classes, against 151 TCRs. We then present an interpretable Bayesian model, called BATMAN, that can predict the set of peptides that activates a TCR. We also developed an active learning version of BATMAN, which can efficiently learn the binding profile of a novel TCR by selecting an informative yet small number of peptides to assay. When validated on our database, BATMAN outperforms existing methods and reveals important biochemical predictors of TCR-peptide interactions. Finally, we demonstrate the broad applicability of BATMAN, including for predicting off-target effects for TCR-based therapies and polyclonal T cell responses.

## Introduction

A single T cell receptor (TCR) can recognize a variety of peptides, a property known as TCR cross-reactivity [1, 2]. Predicting which peptides a TCR cross-reacts to is critical for numerous applications, including predicting viral escape [3], cancer neoantigen immunogenicity [4], autoimmunity [2, 5], and off-target toxicity of T cell-based therapies [6, 7]. However, predicting interactions among TCRs, peptides, and major histocompatibility complexes (TCR-pMHCs) remains challenging [8–11] due to: (a) limited TCR cross-reactivity assay data, (b) few experimentally validated negative examples [12], which are important for model discrimination (Figure 1A), and (c) limited number of available ground-truth TCR-pMHC structures [13]. Most existing computational methods are designed to cluster different TCRs that bind the same peptide [8, 14]. But the opposite task — predicting peptides that bind a given TCR — remains outstanding [9, 11, 15, 16]. This is largely due to the sensitivity required to discriminate among single amino acid (AA) mutants [11, 17] of a TCR’s known index peptide, i.e., the peptide to which the TCR was identified to strongly bind. To address this challenge, we offer both a comprehensive experimental mutational scan database of TCR-pMHC binding, and a method that can predict how peptide mutations affect TCR activation. Additionally, we also developed an active learning method to minimize the number of TCR-pMHC experiments that need to be performed to learn the peptide cross-reactivity of a novel TCR, thereby reducing costs associated with performing TCR-pMHC assays.

**Figure 1.**
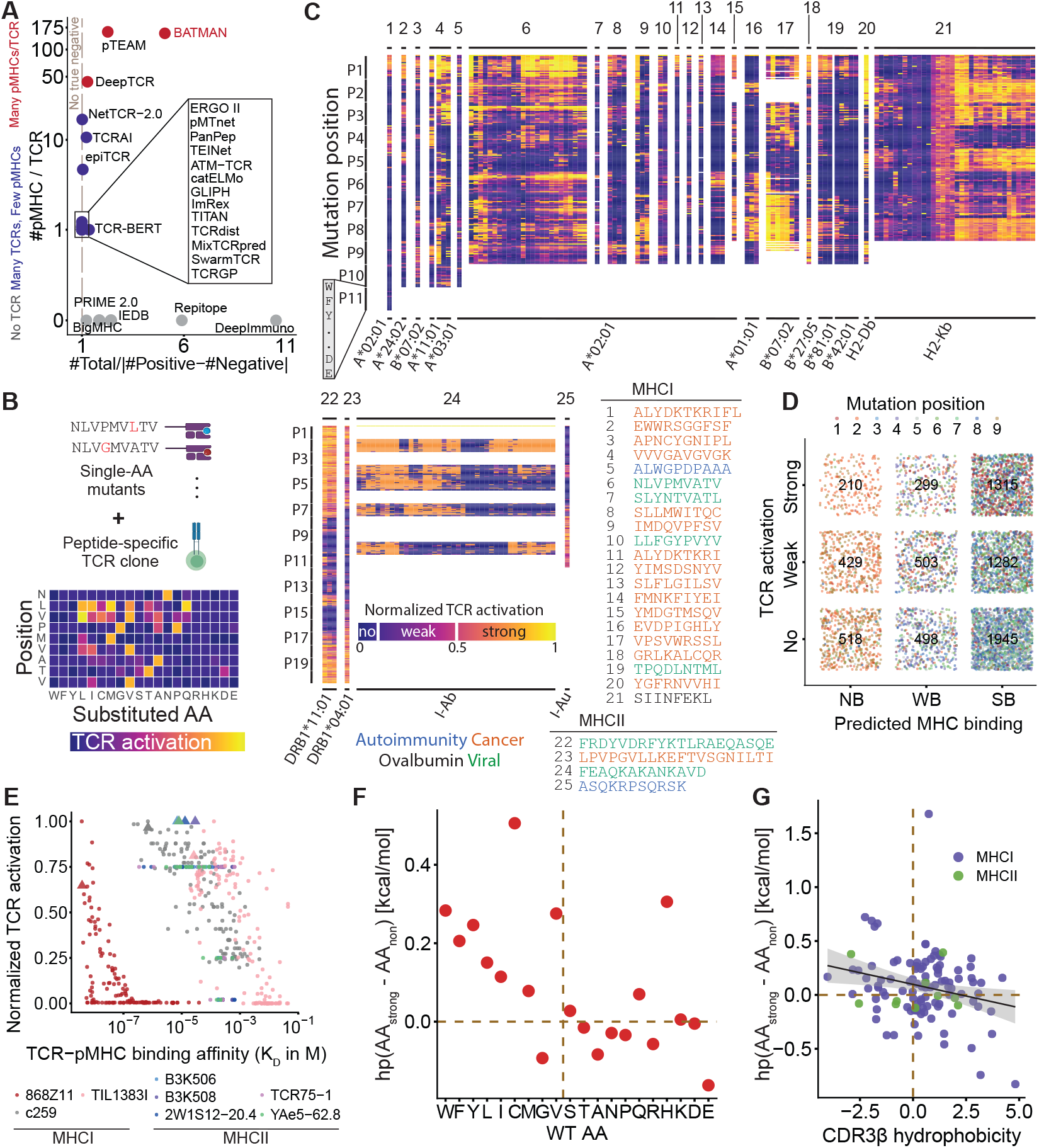
The BATCAVE database reveals important insights into TCR-pMHC interactions. **A**. Training dataset summary metrics for the number of pMHCs tested per TCR and the class balance of experimentally verified TCR-pMHC interactions, shown for selected TCR-pMHC interaction prediction methods. **B**. Schematic of a mutational scan assay, reporting activation of a TCR clone against all single-AA mutants of its index (here, NLVPMVATV). **C**. Curated BATCAVE mutational scan database for TCR activation, with each column corresponding to a TCR clone, grouped by their index peptide (indicated above each column) and recognized MHC (below), and each row corresponding to the substituted AA at a specific position, ordered according to their interfacial hydrophobicity. **D**. Dependency of mutant TCR activation on NetMHCPan-predicted pMHC binding, shown for all HLA-A*02:01-restricted 9-mer binding TCRs in BATCAVE, with the number of mutants (single dot) in each of the nine sub categories (squares) indicated. **E**. Dependency of TCR activation on TCR-pMHC binding, with dots representing TCR-pMHC pairs and TCRs respectively (index peptides indicated by triangles). Hydrophobicity difference between allowed and disallowed AA substitutions depend on WT peptide AA (**F**, dots representing average over all occurrences of a WT AA) and CDR3*β* hydrophobicity (**G**, dots representing TCRs).

### BATCAVE: a database of TCR-specific peptide mutational scans reveals TCR-pMHC interaction rules

To address the lack of TCR cross-reactivity training sets with balanced positive and negative experimentally-validated TCR-pMHC pairs, we curated a database of continuous-valued TCR activation data by pMHCs from mutational scan assays (hereafter referred to as TCR activation data; Figure 1B,C). This database, termed BATCAVE (“**B**enchmark for **A**ctivation of **T**-cells with **C**ross-reactive **Av**idity for **E**pitopes), includes (1) 35 fully-sequenced CD8^+^ and 43 (7 fully-sequenced) CD4^+^ mouse TCR clones, and (2) 69 (59 fully-sequenced) CD8^+^ and 4 fully-sequenced CD4^+^ human TCR clones (Extended Data Fig 1). Together, TCRs in the BATCAVE recognize a total of 25 unique index peptides that are of length 8 to 20 (*L* ∈ [8, 20]) AAs presented by 11 class-I MHCs and 4 class-II MHCs molecules, and which are involved in cancer, viral infection or autoimmunity. For 101 of 151 TCRs, BATCAVE records the raw activation of all possible *L* × 19 single-AA mutant peptides, and for the rest of TCRs, the corresponding data for all mutant peptides with mutations only in non-MHC-anchor positions (Figure 1C). To provide a consistent activation scale across diverse TCR-pMHC avidities and experimentally measured variables (e.g., IFN*γ* concentration and NFAT-GFP fluorescence intensity), we normalized the raw activation data to the maximum activation for each TCR. In addition to the TCR activation data, we collected matched TCR-pMHC dissociation constants of all single-AA mutant peptides for 3 MHCI-restricted TCRs (868Z11, c259, TIL1383I), and 5 MHCII-restricted TCRs (all specific for the H2-IAb-bound ‘3K’ antigen FEAQKAKANKAVD). Overall, we achieved (1) full coverage of the single-AA mutant antigenic space for over 100 TCRs, and (2) balanced numbers of high-confidence, true positive (TCR activated) and true negative (TCR inactive) data for each TCR, both unprecedented among existing datasets (Figure 1A).

BATCAVE showed that single AA changes of the index peptide can result in both loss and gain of TCR activation over orders of magnitude (Figure 1C, Extended Data Fig 2), and it recorded how different mutants of the same index peptide activate different TCRs [18–20], allowing us to distill broad, interpretable biochemical and cellular features of TCR-pMHC interactions. First, we analyzed how pMHC binding affects TCR activation. Since different MHC alleles have different peptide-binding anchor residues, we first focused on the most prevalent MHC allele in our database, HLA-A*02:01 (43 TCRs, 11 index peptides). We selected the 41 TCRs recognizing nine distinct 9-AA-long index peptides with all peptide positions mutagenized, and we used pMHC binding level prediction from NetMHCPan4.0 [21] to classify pMHC interaction as ‘no’ [> 2% rank], ‘weak’ [0.5-2% rank], or ‘strong’ [< 0.5% rank] binders. For ease of interpretation, we also discretized continuous TCR activation values into three levels (Figure 1C): ‘strong’ [≥ 50% of maximum], ‘weak’ [10-50%] or ‘no’ [<10%] activation. We found that the fraction of TCR strong-activating mutants increased from 0.18 for MHC non-binders, to 0.23 (weak-binders) and 0.29 (strong-binders) with predicted MHC binding. However, mutant peptides were distributed over all 3 ×3 = 9 possible paired (MHC-binding level × TCR activation level) categories (Figure 1D), with similar observations for other MHCIs (Extended Data Fig 3). This suggests that MHC binding alone cannot be used to predict TCR activation and demonstrates the need for TCR activation and peptide immunogenicity prediction models beyond pMHC affinity prediction.

Second, we utilized BATCAVE to deduce the relationship between TCR-pMHC binding (measured as *K*_*d*_) and TCR activation. This is a classic question in immunology with many features contributing to the relationship between binding and activation, such as pMHC concentration [22], TCR structural changes [23, 24], catch bonds [25, 26], and co-receptors [27–29]. We found a general positive association between TCR-pMHC binding and TCR activation across both MHCI and MHCII TCRs (Figure 1E). However, there are exceptions; for example, for TCR 868Z11, which binds the disulfide-stabilized HLA-A02:01 molecule [30], we found that strong TCR-pMHC binding of many mutants is not sufficient for TCR activation.

Third, we asked how TCR cross-reactivity depends on the wildtype (WT) AAs of the index peptide. Since previous studies have associated hydrophobic peptide residues with increased immunogenicity [31–33], we asked whether WT AA residue hydrophobicity dictates the hydrophobicity of allowed mutations. To quantify this, we calculated the difference between average hydrophobicity (as measured by the Wimley-White interfacial hydrophobicity scale [34]) of TCR non-activating and strong-activating mutated AAs for each WT AA across all TCRs and peptide positions (Figure 1F). Positive differences in average hydrophobicity between strong-activating and non-activating mutated AAs were more frequent when WT AAs were hydrophobic (89%) than for any other class of WT AA (36%), with the mean differences in hydrophobic WT AA and all other AAs being 0.18 and 0.00, respectively. This indicated that peptide positions with non-hydrophobic WT AAs do not have any preference for the mutant AA for TCR activation, while hydrophobic WT AA residues preferentially allowed hydrophobic mutant AAs, likely since the latter are crucial TCR binding residues.

Finally, we studied how the TCR CDR3 chain hydrophobicity dictates the hydrophobicity and number of allowed mutations. For TCRs with available *CDR*3*α* and *CDR*3*β* sequences, we again calculated the difference between average hydrophobicity of TCR non-activating and TCR-activating mutated peptide AAs. Next, we computed the hydrophobicity of the CDR3*αβ* chains by summing across the hydrophobicity of their individual AA residues. We found that hydrophobic (hydrophilic) CDR3*β* chains prefer hydrophobic (hydrophilic) mutant AAs for activation (Figure 1G), confirming a trend previously seen for TCR-pMHC structural analyses [32, 35, 36].

Overall, BATCAVE provides the largest to-date collection of TCR-pMHC interaction data across multiple MHC classes, index peptides, and TCRs and reveals broad as well as TCR-specific insights into properties of TCR-pMHC binding. Together, this database serves as a strong foundation for benchmarking TCR-pMHC prediction methods.

### Existing TCR-pMHC models poorly predict TCR activation by mutant peptides

We first asked how well existing TCR-pMHC models predict TCR activation by mutant peptides, a question we could answer using BATCAVE as a benchmarking dataset. Since many index peptides appear multiple times for different TCRs in BATCAVE, we selected a subset of BATCAVE with each index peptide appearing only once. This selection yielded a diverse and balanced dataset, spanning 18 MHCI-restricted human TCRs specific for unique 9- and 10-AA-long index peptides and 4 MHCII-restricted TCRs specific for index peptides of various lengths (Figure 2A). For TCRs with the same index peptide, we included the one with the least class imbalance.

We benchmarked 15 pre-trained TCR-pMHC models with different machine learning architectures and input data requirements (Figure 2A), selected based on the availability of web-servers or source code. These models are trained on large publicly available databases (e.g., VDJdb [37], McPAS-TCR [38], and single-cell immune repertoire profiling data [39]). While these methods are designed for either classifying TCRs binding to selected immunogenic peptides [8, 14, 40] or predicting peptide immunogenicity [32, 41], our benchmarking involved a distinct problem of classifying which set of peptides activate a given TCR. Similar to previous results on smaller and less diverse datasets [11, 17], these models predicted only marginally better than random on our benchmarking dataset, with the best classification AUCs being from TITAN [42] (mean AUC=0.56 for MHCI-restricted and 0.52 for MHCII-restricted peptides) and ImRex [9] (mean AUC=0.55 for MHCI-restricted peptides). General immunogenicity prediction tools, which do not use TCR sequence information for prediction, overall performed worse, with none included among the top 6 performers. Overall, predicted TCR-pMHC interaction scores from most tested TCR-pMHC models were uncorrelated with true TCR activation values for the mutant peptides (Extended Data Fig 4). Thus, these results demanded development of new TCR-pMHC models for predicting how peptide mutations affect TCR activation.

**Figure 2.**
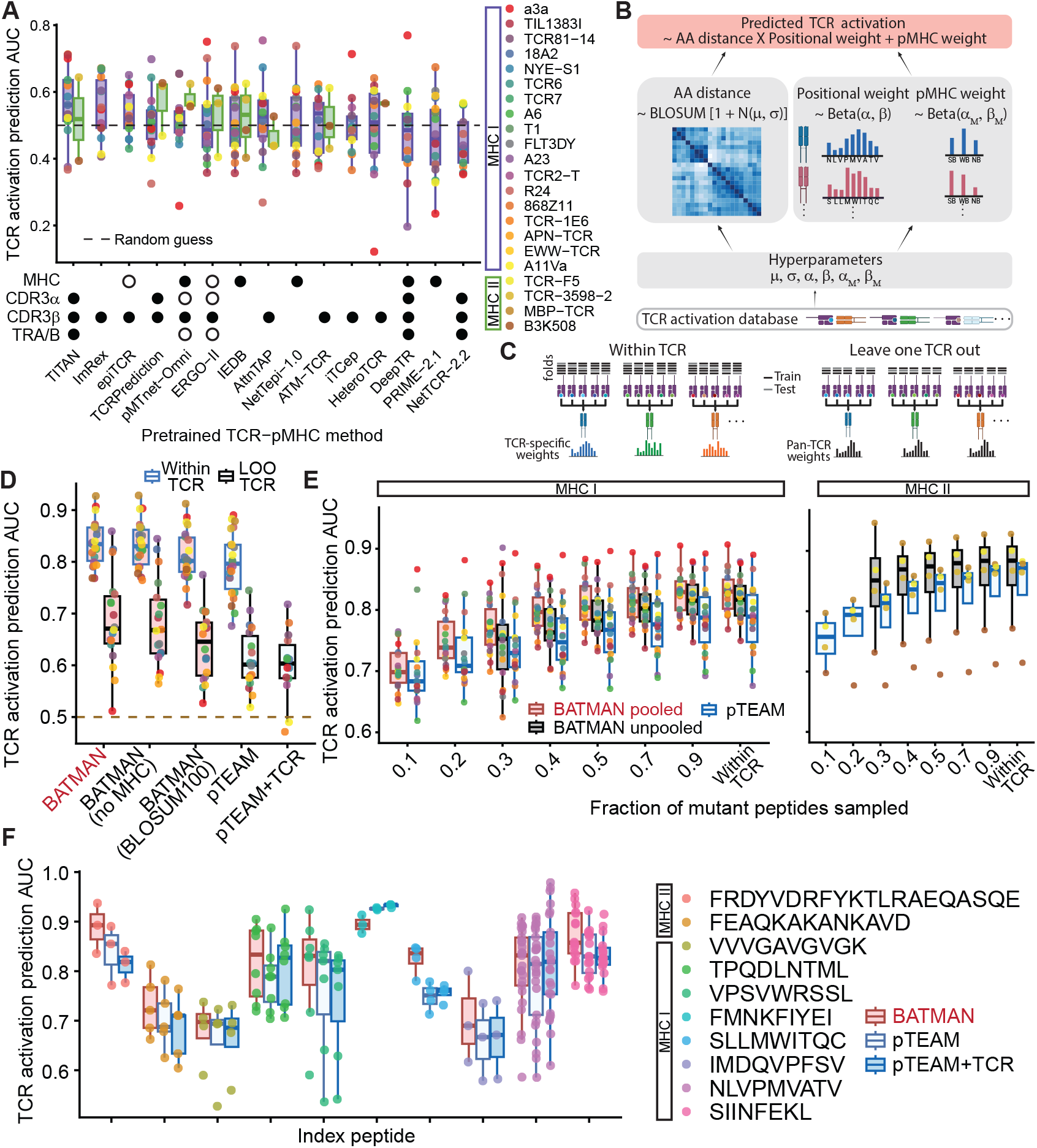
BATMAN outperforms existing TCR-pMHC interaction prediction methods. **A** TCR activation classification area under the curve (AUC) scores for different pre-trained methods, with their respective requirements indicated by dot matrix (filled: mandatory, open: optional, used if available in database); MHC: name of peptide-binding MHC allele; CDR3*α*/*β* : sequence of the TCR CDR3*α*/*β* chain; TRA/B: gene name or sequence of at least one of the TRAB, TRAJ, TRBV, and TRBJ gene name or sequence. Schematics for **B** BATMAN TCR activation prediction method and **C** two validation modes, within-TCR and LOO-TCR. **D**. BATMAN outperforms pTEAM in both within-TCR and LOO-TCR modes. **E**. BATMAN outperforms pTEAM for different numbers of training samples, showing improvement with pooling across TCRs (TCR color scheme same as in A). **F** BATMAN and pTEAM LOO-TCR performance comparison among TCRs specific for a common index peptide.

### BATMAN: a Bayesian inference model provides state-of-the-art prediction of TCR activation by mutant peptides

We present BATMAN — “**B**ayesian Inference of **A**ctivation of **T**CR by **M**utant **An**tigens” — a hierarchical Bayesian model that can predict TCR activation by single-AA mutant peptides based on their distances to the TCR’s index peptide. Our peptide-to-index distance is a product of (a) a learned positional weight profile, specific for individual TCRs, corresponding to the weighted effects of mutated residues at different positions in the sequence, (b) a learned AA substitution distance matrix from the index AA to the mutant AA, and (c) an optional scalar weight, also specific for individual TCRs, corresponding to the pMHC binding category (Figure 2B, Methods). BATMAN can be used for classification and continuous regression tasks for both TCR-specific and cross-TCR activation datasets.

We validated BATMAN over our benchmarking dataset in two modes (Figure 2C):

1. *Within-TCR* mode, where we performed 5-fold cross-validation for each TCR, with the folds stratified by TCR activation category and mutation position. The inferred positional weight profiles and pMHC binding level weights were TCR-specific, while the inferred AA matrix was TCR independent. Even though we inferred TCR-specific parameters, the hierarchical Bayesian framework of BATMAN allowed pooling of training data across TCRs to infer TCR-independent model hyperparameters and the AA matrix (see Methods).
2. *Leave-one-TCR-out (LOO-TCR)* mode, where we trained on data from all TCRs but one, and tested on the held-out TCR. The inferred positional weight profiles, pMHC binding level weights, and inferred AA matrix were all TCR-independent, and learned by training across all TCRs except the held-out one (see Methods).

Since all of the pretrained TCR-pMHC models (Figure 2A) performed close to random in our benchmarking study, we compared BATMAN’s performance to that of *pTEAM* [17], an existing method designed to predict the effect of peptide mutation effects on TCR activation. For a fair comparison, we re-trained *pTEAM* using the same data as BATMAN.

BATMAN outperformed *pTEAM* in both within-TCR (mean AUC=0.836 versus *pTEAM* AUC=0.795) and LOO-TCR (mean AUC=0.687 versus *pTEAM* AUC=0.613) classification (Figure 2D). Critical to achieving BATMAN’s performance was learning the AA distance matrices by pooling training data across TCRs. For example, applying BATMAN with the BLOSUM100 AA matrix, the best-performing conventionally-used distance matrix (Extended Data Figs 5 to 7), dropped the within-TCR AUC to 0.811 (Figure 2D). Extended Data Figs 5 to 7 further highlight the superior performance of BATMAN over 68 conventional AA substitution distance matrices when tested on all 151 TCRs, both in classification and continous regression tasks. BATMAN performance also slightly improved with the addition of pMHC binding level information, with the within-TCR AUC dropping to 0.833 without this input (Figure 2D). *pTEAM* can also incorporate TCR sequence information into its model, but we found no gain in performance with this additional information (Figure 2D), showing its limitations when the training data is sufficiently diverse.

How do BATMAN and *pTEAM* compare when training data becomes limited? We calculated the within-TCR AUCs from both models over random subsamples of previous training folds, maintaining stratification over TCR activation classes and mutation positions. In each case, we also trained within-TCR BATMAN in two ways: (1) pooled across all TCRs like before, and, to investigate effects of pooling across TCRs and the overall training data size, (2) trained and tested individually for each TCR. In this limited-data regime, BATMAN pooled across TCRs consistently performs better than *pTEAM*, with BATMAN achieving close to its 5-fold cross-validation performance with only a random 40% subsample of training data in each fold (7 per mutation position per TCR) (Figure 2E). Pooled BATMAN performed better than unpooled BATMAN, for which MHCI AUC dropped to 0.775 for the above 40% subsampling, though the difference narrowed with more training data (MHCI AUC=0.830 and 0.817 respectively for unpooled and pooled with full within-TCR data; see also Extended Data Fig 7). Unpooled BATMAN also performed comparably to *pTEAM* in the very low training data regime (Figure 2E). Overall, these results highlight the unique ability of BATMAN to improve TCR-pMHC interaction prediction by pooling across TCRs compared to *pTEAM*, which must be trained separately for every TCR.

Finally, we compared BATMAN and *pTEAM* for LOO-TCR tasks where the training data is restricted to TCRs with the same index peptide. For this benchmark, we selected 8 MHCI and 2 MHCII index peptides for which mutational scan data against at least 3 TCRs were found in BATCAVE. BATMAN performed better than *pTEAM* for 9 out of 10 index peptides tested (Figure 2F), with their performances becoming comparable only for index peptides with a large number of TCRs tested (e.g., TPQDLNTML (n=8), NLVPMVATV (n=25) and SIINFEKL (n=16)). These benchmarking results collectively establish BATMAN as the state-of-the-art predictor of TCR activation by mutant peptides across diverse TCR-pMHC training data contexts.

### Inferred BATMAN parameters capture interpretable features of TCR-pMHC interactions

BATMAN inferred positional weights were consistent across different AA matrices (Extended Data Fig 8), indicating that they might correspond to key TCR-pMHC interaction features. Thus, we asked what the inferred positional weight profiles and AA substitution matrices of BATMAN can reveal about TCR-pMHC interaction rules.

First, we inferred a pan-TCR positional weight profile and AA matrix by training BATMAN over all TCRs binding to 9-AA-long peptides in our benchmarking dataset of Figure 2A. The learned pan-TCR positional weights (Figure 3A) peaked near the middle of the peptide chain, reflecting the fact that central AA residues more directly affect TCR binding compared to flanking MHC-anchor residues [19, 41, 43–46] (see also Extended Data Fig 2).

**Figure 3.**
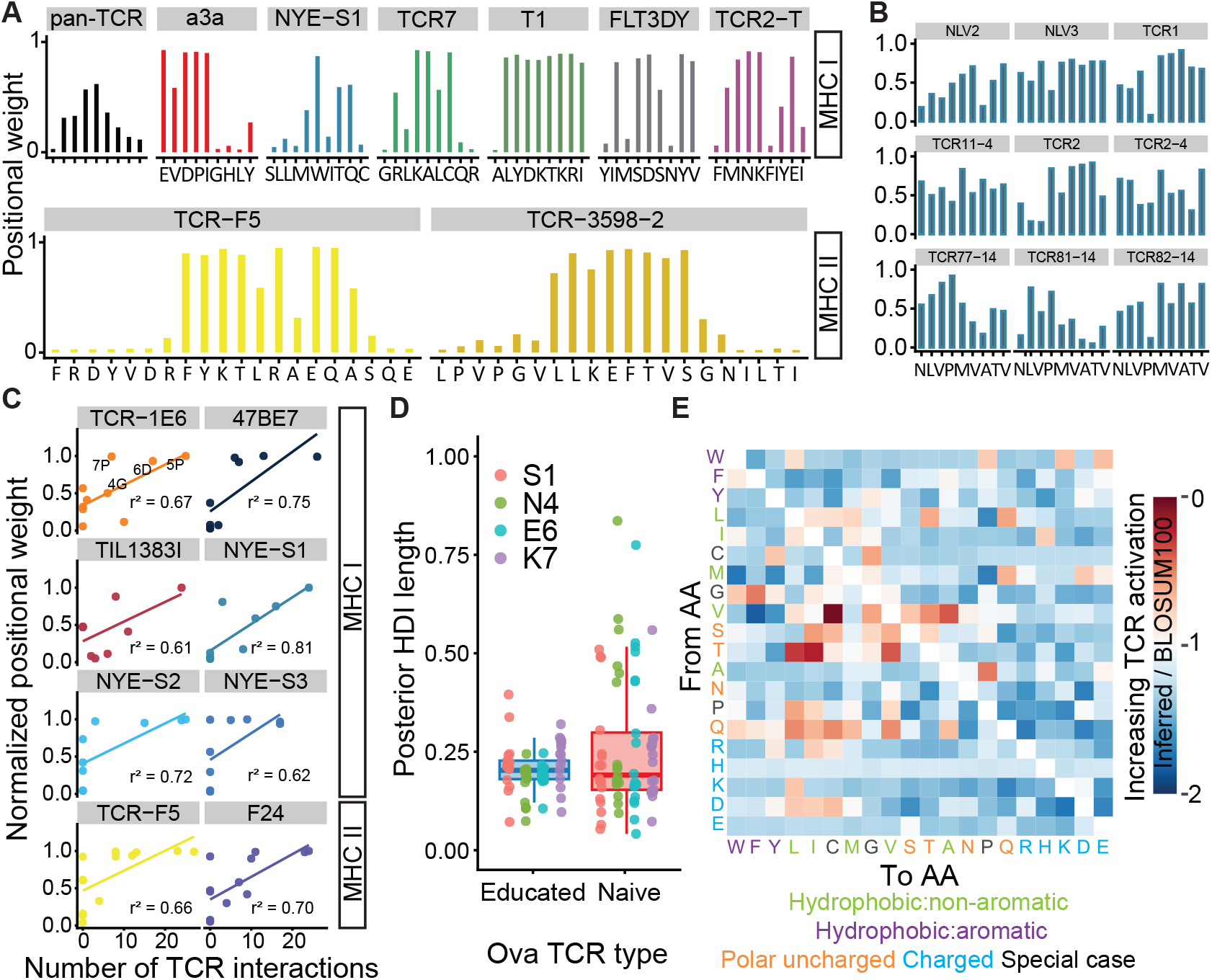
Inferred BATMAN parameters capture interpretable biochemical features of TCR-pMHC interactions. **A**. pan-TCR positional weights inferred from all 9-mer-binding TCRs in benchmark dataset (first panle), and examples of TCR-specific positional weights of selected TCRs with different index peptides (remainder of panels). **B**. TCR-specific positional weights of nine different NLV-specific TCRs, showing important peptide residues for TCR binding. **C**. Inferred positional weights correlate the number of interactions of peptide with TCR chains (derived from TCR-pMHC structure). To make different weight profiles comparable across TCR panels, we normalized positional weights by their maximum over all positions for each weight profile. 4 TCR-binding peptide residues are highlighted for TCR-1E6. **D**. 90% high-density intervals (HDIs) of positional weight posterior distributions for non-MHC-anchor positions are wider for low-avidity ‘naive’ SIINFEKL-specific TCRs than SIINFEKL-expanded ‘educated’ ones. **E**. Example ratio of inferred matrix elements to BLOSUM100, the best performing conventional AA distance matrix, for within-TCR classification.

Second, we pooled across TCRs but inferred TCR-specific weight profiles, which showed that while central residues are indeed often highly weighted, there can be significant variability of binding motifs for individual TCRs (Figure 3A, Extended Data Fig 8), suggesting that TCR cross-reactivity constitutes a diverse spectrum. Some TCRs (e.g., a3a, T1) are activated when very specific AAs exist at certain positions (e.g., P2-P9 for T1, P1-P5 for a3a), and by almost all AAs at other positions (P1 and P6-P8, respectively). These TCRs have almost binary positional weight profiles and resulted in better within-TCR classification performance (Figure 2D). Similarly, positional weights could reveal how different TCRs that bind the same index peptide use different binding motifs. We trained BATMAN by pooling over all 25 BATCAVE TCRs specific for the NLV peptide and indeed found heterogeneity in positional weights across TCRs (Figure 3B). For example, the M5 residue is important for binding to most TCRs, while N1 and A7 residues show high variation in their importance across TCRs.

Third, we asked whether BATMAN can identify the exact TCR epitopes from mutational scans of longer MHCII-bound peptides, which would be important for T cell epitope discovery from peptidome-wide tiling scans (e.g. [47]). To test this, we investigated BATMAN weights for the CD4^+^ TCRs TCR-F5 and TCR-3598-2 in the TScan-II dataset [47] performing mutational scans across all positions in 20-AA-long peptides. The inferred positional weights (Figure 3A) peaked at peptide positions included in the experimentally-verified central TCR-binding epitopes, FRDYVDR**FYKTLRAEQA**SQE [48] and LPVPGV**LLKEFTVSG**NILTI [47]. This suggests BATMAN weights can reveal the exact TCR epitopes from longer peptide tiles with many TCR-irrelevant positions mutagenized.

Fourth, to study the correspondence between BATMAN positional weights and TCR-pMHC 3D structures quantitatively, we assessed the interactions between peptide residues and TCR chains for 6 MHCI and 2 MHCII TCRs in BATCAVE with available ground-truth crystallographic structures of their corresponding TCR-pMHC complexes in the Protein Data Bank (PDB). BATMAN positional weights correlated strongly (*r*^2^ between 0.61-0.81) with the number of interactions between peptide residues and TCR chains (Figure 3C). For example, for the highly cross-reactive [2] autoimmune TCR 1E6 ([49], PDB ID: 3UTT) binding the insulin peptide ALW**GPD**PAAA, positional weights are large for the solvent exposed GPD bulge of the peptide, over which the TCR binds [49]. Similar correspondence between large inferred positional weights and important TCR recognition motifs on peptides, characterized by a large number of TCR interactions, were seen for the other TCRs (Figure 3C). This suggests that BATMAN weights encapsulate key structural features of TCR-pMHC complexes that affect TCR-peptide binding.

Fifth, we investigated additional features of TCR-pMHC interactions revealed by the inferred positional weight profiles. As a test case, we used a dataset spanning a wide range of cross-reactivity, consisting of both low-avidity ‘naive’ (broadly cross-reactive) and antigen-expanded ‘educated’ (specific) TCRs binding to the SIINFEKL peptide [18]. We trained BATMAN by pooling over all TCRs and calculated the width of the posterior distribution of the inferred positional weights for each peptide residue which contains 90% of the density. To eliminate differences among TCRs due to their different pMHC binding requirements, we focused only on the non-MHC-anchor positions (P1, P4, P6, P7) of the SIINFEKL peptide. We found that naive TCRs with broader cross-reactivity profile have broader posterior distributions of inferred weights (Figure 3D), suggesting that the weight profiles capture essential features of cross-reactivity.

Finally, the BATMAN AA distance matrix (Figure 3E) revealed three biochemical features of TCR-pMHC interactions. First, large changes in TCR activation correspond to non-aromatic to aromatic AA substitutions (e.g., valine and glycine to phenylalanine; methionine to tryptophan and tyrosine) affecting side-chain interactions [31, 32, 41]. Second, swapping in hydrophobic isoleucine and leucine residues for non-hydrophobic residues overall increases TCR activation, in line with these residues considered to increase immunogenicity [31–33] and our earlier observation in BATCAVE data (Figure 1F) that strong-activating mutant AA residues are more hydrophobic than non-activating residues for highly hydrophobic WT AAs. Third, while previous works have reported both gain [50] and loss [51] of TCR activation with peptide cysteine residue substitutions, we found an increase in TCR activation for valine, glutamine and serine to cysteine substitution while there was a decrease in activation for phenylalanine and tyrosine to cysteine substitutions (Figure 3E), reflecting the high context-dependence of such substitutions.

Overall, the interpretable parameters learned by BATMAN capture a host of biological features that reveal the nature of TCR-pMHC interactions.

### Active learning with BATMAN boosts performance for novel TCRs

Designing cross-reactivity assays for a novel TCR is challenging because the number of possible peptides to test increases exponentially with the number of mutations. Can we identify a small set of informative peptides within this space, whose binding information can be used to predict TCR activation to the many other peptides not assayed?

We developed a BATMAN-guided active learning framework (called ‘BATMAN-AL’), which iteratively selects a small number of mutant peptides that provide the maximum information about the cross-reactivity of a given TCR. Intuitively, these peptides should (a) be diverse, so as to adequately cover the peptide space, and (b) lie near current decision boundaries, thus helping classifiers better delineate between activation classes. Specifically, in each experimental round, the TCR binding of a set of mutant peptides is determined experimentally, and then the binding results are fed-back to update the model, which subsequently picks the next set of maximally informative peptides to test in the next round, until predictive performance converges. Computationally, it may be advantageous to select only one peptide per round, so that the model can use binding information from this peptide to better select the next peptide; however, this may be experimentally prohibitive because of constraints on the total number of rounds a TCR can be assayed, thus requiring the selection of multiple peptides per round. To address these requirements, we developed AL strategies to maximize classification performance boost of each AL round by sampling the most informative and diverse set of peptides in each round (Extended Data Fig 9).

BATMAN-AL achieved a superior classification performance with mean AUC of 0.786, only slightly lower than the AUC for within-TCR task (which used 144 training peptides), using only ~ 45 total peptides per TCR selected over 5 rounds (Figure 4A). Even if constrained to only select 9 total peptides in a single round, BATMAN-AL achieved a mean AUC of 0.730, outperforming the leave-one-TCR-out mean AUC of 0.687 (Figure 4A), which suggests that even a minimal amount of active learning can boost performance. BATMAN-AL also outperformed random active learning (BATMAN-RL; Figure 4A), where training peptides are chosen randomly. Furthermore, BATMAN-AL demonstrated much less variance across TCRs compared to BATMAN under the LOO-TCR mode, and the variance was comparable to within-TCR levels after only 5 rounds, demonstrating how small, carefully chosen training set sizes can boost robustness across TCRs.

**Figure 4.**
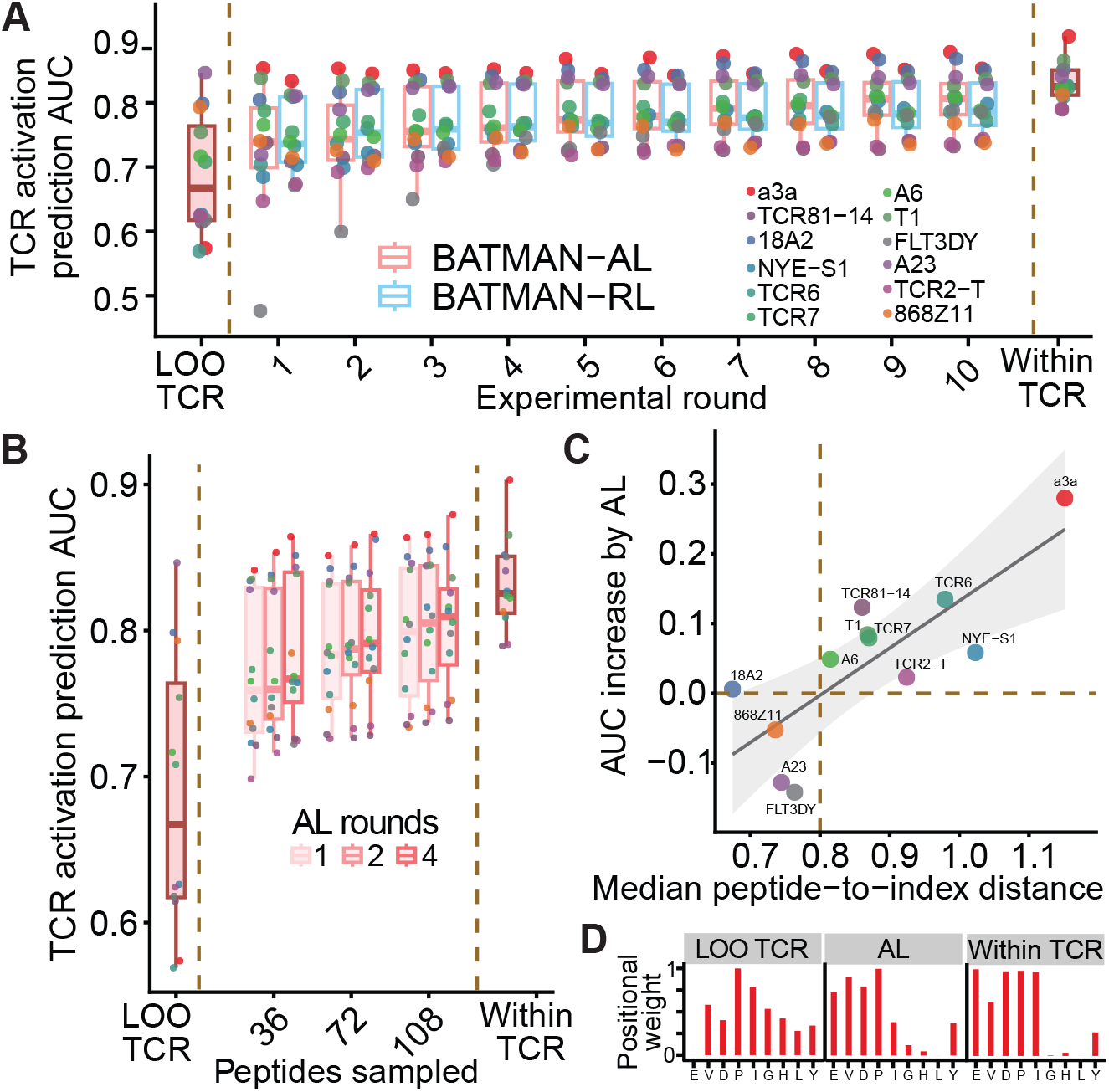
Active learning (AL) with BATMAN predicts TCR-specific activation with limited training samples. **A**. Average AUCs for BATMAN with active (AL) and random learning (RL). At each learning step (Experimental round, x-axis), 9 peptides are sampled. **B**. BATMAN-AL performance when the same total number of peptides are sampled over different numbers of AL rounds. For performance comparison in panels **A** and **B**, leave-one-TCR-out and within-TCR AUCs are shown. **C**. Larger median peptide-to-index distances are associated with higher AL performance boost with BATMAN. **D**. After only one AL round with 9 peptides, the positional weight profile of the a3a TCR approximates the within-TCR weight profile.

We next explored the performance of BATMAN-AL under various constraints on the total number of peptides to test and number of experimental rounds. For example, if a fixed number of peptides are tested (say, *N*_*p*_), is it best to select all *N*_*p*_ of them in a single round, or might it be better to select *N*_*p*_*/*4 peptides per round over 4 rounds? In general, we found that performing more rounds of experiments for a fixed total number of peptides improves predictive performance (Figure 4B; e.g., for *N*_*p*_ = 72, mean AUC increases from 0.787 in single round to 0.798 for 4 rounds), although for fewer total peptides (e.g., *N*_*p*_ = 36), a single active learning round is sufficient. Testing more peptides, unsurprisingly, improves performance, although if a large number of peptides can be tested (e.g., *N*_*p*_ = 108), a single active learning round is again sufficient to achieve within-TCR-level performance.

What determines how much BATMAN will benefit from active learning for a novel TCR? We found that larger median peptide-to-index distances of all single-AA mutants of its index peptide (calculated using the pan-TCR weight profile and AA substitution matrix) correlated with performance boost with one AL round with 9 peptides (Figure 4C). One reason for this could be that larger median distances indicate a broad range of effects for single-AA mutants (Extended Data Fig 9A), and thus better separability between classes that can be learned by AL. Furthermore, TCRs with smaller median peptide-to-index distances often corresponded to narrower within-TCR-activation-class distance distributions (e.g., among strong-activating mutants of TCRs A23 and 18A2 in Extended Data Fig 9A), and so already better LOO-TCR classification performance than other TCRs, gaining only a slight additional performance boost by AL (Figure 4A). For example, TCR a3a, having the largest median peptide-to-index distance, demonstrated the most improvement with AL, with its AUC approaching within-TCR levels after one AL round using 9 peptides (Figure 4A). Comparison of a3a positional weights profiles (Figure 4D) showed that after the single AL round, the weight profile indeed closely matched the within-TCR profile.

In summary, BATMAN-AL can efficiently learn the binding profile of novel TCRs from only a handful of example and can provide custom testing strategies based on experimental constraints.

### BATMAN TCR activation predictions generalize to multiple peptide mutations

While BATCAVE records effects of single-AA mutations on activation of individual TCRs, we asked whether BATMAN predictions can be generalized beyond single-AA mutants. For this benchmark, we collected multi-AA mutant TCR activation datasets for the BATCAVE TCRs for which single-AA mutants were evaluated by the same assay in the same source publication (n=6 TCRs). While BATMAN benchmarked on single-AA mutants of unique index peptides showed significant improvement with pooling across TCRs (Figure 2E), here we used unpooled BATMAN because (1) the training set is biased in the number of TCRs per index peptide (3 MAGE-A3-specific TCRs, but only one specific TCR for the 3 other index peptides), and (2) the training set consists of full single-AA data while the test set consists of all available multi-AA mutant data. Thus, for each TCR, we trained BATMAN in unpooled within-TCR mode over the full single-AA data of individual TCRs, and inferred the TCR-specific positional weights and AA matrix as before. We defined the peptide-to-index distance of multi-AA mutants by summing over the product of positional weight and AA substitution distance for all mutational positions. We then used the peptide-to-index distances to rank multi-AA mutants and investigated whether this ranking predicts TCR activation.

We found that BATMAN could identify strong TCR activating mutants at all Hamming distances (Figure 5A). For example, among 32 Hamming distance 8 mutants for TCR c259 (index peptide SLLMWITQC), the sole binder NVSL**W**LSAV [52] was correctly ranked as top. As another motivating example, we considered the a3a TCR, which was designed to target the cancer-testis antigen MAGE-A3 (EVDPIGHLY). However, it resulted in fatal off-target cross-reactivity to muscle antigen Titin (**E**S**DPI**VAQ**Y**), differing from MAGE-A3 by 4 AA mutations [53, 54]. We found that BATMAN, trained on only single-AA mutants, ranked Titin 6th among 29 4-AA mutants and 8th among all 111 multi-AA mutants in our a3a dataset. These results indicate that a complete single-AA mutational scan can generalize to some extent to multi-AA mutated cross-reactive targets using BATMAN.

Can BATMAN-AL better identify TCR-activating multi-AA mutants from only single-AA mutant training? We tested this using the a3a TCR and found that the peptide-to-index distances of BATMAN-AL, using only 9 single-AA mutants, performed better than pan-TCR distances in classifying multi-AA mutant activation, with the rank of Titin improving from 41st to 16th among 111 multi-AA mutants in our a3a dataset (Figure 5B). These results suggest that BATMAN and BATMAN-AL predictions can identify cross-reactive targets for a novel TCR beyond those in the 1-Hamming space.

**Figure 5.**
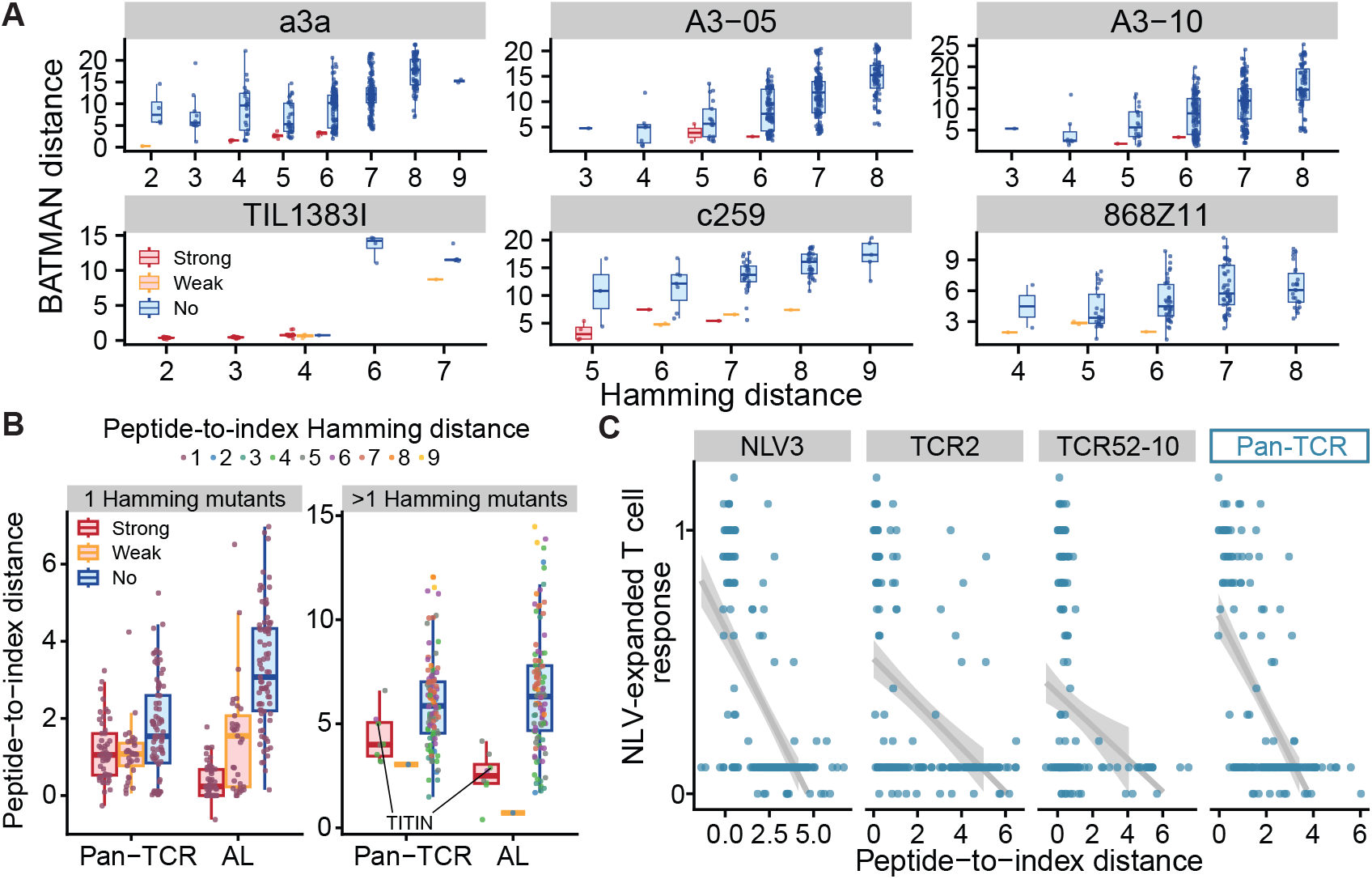
BATMAN predictions generalize to multiple-AA mutations and polyclonal T cell response. **A**. BATMAN peptide-to-index distances from single-AA mutant training separates multi-AA mutants according to their TCR activation. **B**. BATMAN-AL with only 9 peptides for the a3a TCR allows improved TCR activation prediction of both 1 Hamming and >1 Hamming mutant peptides. **C**. NLV-expanded polyclonal T cell repertoire response against NLV mutant peptides correlates with the response of individual NLV-specific TCRs to various degrees (shown are results from 3 random example TCRs), whereas Pan-TCR response, predicted from pooling across all 25 NLV-specific BATCAVE TCRs, correlating more strongly.

### BATMAN predicts polyclonal T cell response

While we have thus focused on predicting activation of individual TCR clones against index peptide mutations, in some situations (e.g., predicting neoantigen immunogenicity, viral escape mutants), we are interested in the collective response of T cell clones present in a polyclonal repertoire with different frequencies. For example, if a viral peptide elicits strong response from the peptide-specific polyclonal T cell repertoire, can BATMAN predict which mutants of that peptide could fail to elicit response from the repertoire, and thus may lead to viral escape? For the Human Cytomegalovirus peptide (NLVPMVATV), BATCAVE contained full mutational scan training data for 25 individual TCRs. For testing BATMAN predictions on polyclonal T cell response, we collected the TScan assay dataset measuring continuous-valued, NLV-expanded polyclonal TCR repertoire response towards all single-AA NLV mutants individually [55].

How well can individual NLV-specific TCR clone activation data in BATCAVE predict polyclonal NLV-specific TCR response? To test this, we trained BATMAN in within-TCR mode, over the full single-AA mutant data for individual TCR clones and inferred the AA matrix and TCR-specific positional weights as before. We then used the resulting weights to calculate TCR-specific peptide-to-index distances for all NLV mutants, and calculated their correlations with the polyclonal T cell response. Different NLV-specific TCRs recognize different NLV mutants [17, 55] and thus, are representative of the full polyclonal response to various degrees. This was reflected in the variable correlations of the peptide-to-index distances from individual TCRs with the polyclonal response (e.g., r=0.63 for NLV3 but r=0.47 for TCR2 and r=0.25 for TCR52-10; Figure 5C), indicating that individual NLV-specific TCR clone activation data to NLV mutants cannot be reliably used to predict polyclonal peptide-specific T cell responses.

In contrast, when training BATMAN in pan-TCR mode on all NLV-specific TCR data in BATCAVE, pan-TCR predictions correlated strongly (r=0.61; Figure 5C) to the polyclonal response. This indicates that pooling across TCRs is again critical, as the pan-TCR prediction found from pooling across individual TCRs statistically resembles the polyclonal response more accurately than individual TCRs. These results suggest that BATMAN provides useful predictions about polyclonal T cell response, peptide immunogenicity, and potential viral escape mutants.

## Discussion

We curated BATCAVE, a comprehensive database of experimentally validated positive and negative TCR-pMHC interactions from mutational scan and TCR-pMHC binding affinity assays. BATCAVE contains raw and normalized TCR activation data for both single- and multi-AA peptide mutations, and for both MHCI-restricted and MHCII-restricted TCRs. In addition, BATCAVE contains full TCR sequence information and pMHC binding affinities when available. Using this database, we unraveled the nature of TCR cross-reactivity, including (1) the relationship between pMHC binding and TCR activation; (2) the relationship between TCR-pMHC binding and TCR activation; and (3) insights about how biochemical features (e.g., hydrophobicity) of the TCR’s wildtype AA affects TCR binding.

Despite these advances, there are some biases in BATCAVE that require careful interpretation. First, many datasets included used *in vitro* TCR-pMHC assays with, e.g., UV-mediated peptide exchange for MHC loading (see supplementary notes), and thus the TCR activation level by such pMHCs (Figure 1D) might be different than *in vivo* settings. Second, most BATCAVE TCRs belong to peripheral cytotoxic T cells, which survived thymic selection [56]. This thymic selection bias on TCR repertoire [57–62] may affect, for example, the relationship between pMHC binding and TCR activation, or between TCR activation and TCR-pMHC binding affinity. Third, recorded in BATCAVE as a single number, T cell activation for a given TCR-pMHC pair can be heterogeneous, depending on, e.g., expression of co-signaling molecules [63, 64] (e.g., CD5, CD8, and CD4). Notwithstanding these caveats, we believe this database will provide a strong benchmark for evaluating future methods and for uncovering further principles of TCR cross-reactivity.

Using this database, we then developed a computational method, called BATMAN, that predicts TCR activation of peptides based on their distances to the TCR’s index peptide. BATMAN achieved state-of-the-art performance for predicting cross-reactivity across diverse TCRs and index peptides present in the BATCAVE database. Crucial towards BATMAN’s performance were model-inferred positional weights and AA substitution matrices. In comparison, simple distance functions based on mismatch rules (e.g., Hamming distance) with equal weights assigned to each position proved far from sufficient to accurately characterize cross-reactivity. Furthermore, using the learned weights and substitution matrices allowed us to project cross-reactivity outside the single-AA-mutational space, which we demonstrated by accurately ranking immunogenic peptides of clinically relevant TCRs. In addition, the parameters learned by BATMAN revealed striking variation in weight profiles for different TCRs, which was also supported by structural analyses. We then developed an active learning version of BATMAN, which helps BATMAN generalize to novel TCRs. This version provides an efficient way to sample from the prohibitively large antigenic space by iteratively selecting peptides to assay that provide the best improvement of novel TCR activation prediction accuracy. Overall, BATMAN fills a hitherto unoccupied niche of TCR-pMHC prediction methods by accurately discriminating between small differences in peptide sequences, which we show existing methods fail to predict, but which are essential for understanding neoantigen immunogenicity and off-target effects of TCR-based therapies.

Finally, there are several directions of future work and corresponding applications enabled by this study. First, BATMAN could be improved by incorporating TCR sequence information into the model and by training on datasets from other types of experimental TCR cross-reactivity assays [65] (e.g., yeast display library enrichment [5, 66], T-Scan [55], and SABR [67]), which sample more comprehensively outside the one AA-mutational scan space. Second, insights from our database and method could be used to predict how TCR sequences may determine the relationship between TCR binding affinity and cross-reactivity, and to design high-affinity TCRs with limited off-target cross-reactivity, which has numerous clinical applications.

## Author contributions

AB, DJP, SN and HVM conceptualized the work; AB developed the software; AB and DJP designed the model; AB and CW implemented the user interface with help from SRC; AB and PB curated the database; AB conducted all formal analyses except Protein Data Bank structure analyses (AA and HVM); AB, SN, and HVM wrote the original draft; all authors reviewed and edited the final draft; SN and HVM supervised the work.

## Competing interests

The authors declare no competing interests.

## Acknowledgments

We thank our lab members and the anonymous referees for discussion and feedback on method development and figure design, Paul G. Thomas and Zachary Sethna for discussion on TCR datasets, Anastasia Troshina and Vasilisa A. Kovaleva for designing the BATMAN logo, and all authors who shared their datasets with us. TCR-pMHC schematics in Figure 1B, Figure 2B,C were created with BioRender (https://www.biorender.com/).

## Funding

The research was supported by the Simons Center for Quantitative Biology at Cold Spring Harbor Laboratory; US National Institutes of Health Grants S10OD028632 and 1R01AI167862; and the Simons Pivot Fellowship. This work was discussed in part at the Aspen Center for Physics, which is supported by National Science Foundation grant PHY-2210452. The funders had no role in the template design or decision to publish.

## Methods

### TCR activation dataset collection and processing

We collected continuous TCR-pMHC datasets for complete single-AA mutational scans from all publications containing raw datasets (n=21). To normalize datasets across publications, we scaled TCR activation values by the maximum activation value over all recorded peptides tested against that TCR. The only exceptions to this normalization scheme were for experiments where the TCR activation measurements were on a logarithmic scale (e.g., EC50 values), in which case we used the logarithm of the TCR activation values and linearly transformed them to map to the [0,1] interval. Following previous works [17], we discretized the normalized TCR activation values to 3 ordered levels for downstream classification tasks: no activation (*a*_*no*_ ∈ [0, 0.1)), weak activation (*a*_*weak*_ ∈ [0.1, 0.5)), and strong activation (*a*_*strong*_ ∈ [0.5, 1]). For regression tasks, we directly used the normalized TCR activation values. More technical TCR-specific notes on data collection and processing, as well as links to source publications, can be found in the Supplementary Notes. A number of publications (see Supplementary Materials for citations) contained further mutational scan experiments relevant for our database, but the associated raw datasets were not readily available to us.

To select the benchmark subset of TCRs (e.g., for Figure 2A,D) among all TCRs specific for a common index peptide, we selected the one with the highest balance of TCR activation classes. For this purpose, for each TCR, we calculated the entropy of the frequency distribution of the TCR activation classes, as follows

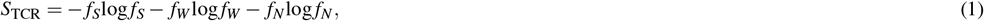

with *f*_*S*_, *f*_*W*_, *f*_*N*_ being the fraction of strong-, weak-, and non-activating single-AA mutant peptides respectively. For each unique index peptide, we selected the TCR with the highest *S*_TCR_. The only exception to this rule was for the index peptide FEAQKAKANKAVD; while all other index peptides had TCR sequences available for either all or none of their specific TCRs, only a subset of FEAQKAKANKAVD-specific TCRs had available sequence. So, for this peptide, we selected the TCR B3K508, with the second highest *S*_TCR_ among FEAQKAKANKAVD-specific TCRs, since the corresponding TCR sequence, necessary for many TCR-pMHC model predictions in Figure 2A, was available.

### Web application for visualizing TCR-pMHC interactions from our database

TCR-pMHC interactions from our database (Figure 1B) are visualized via the web application at https://batman.cshl.edu/. All interactive plots are deployed as a RShiny application, using *ShinyDashboard* (v. 0.7.2). The scatter plot displaying peptide clustering based on index-to-mutant distance was generated via *ggplot2* (v. 2_3.4.4) and rendered using *plotly* (v. 4.10.3). The heatmap presenting normalized peptide activation per index peptide was generated with *InteractiveComplexHeatmap* (v. 1.8.0). The Alluvium plot visualizing the binding of index and mutated peptides to TCRs was generated with *ggplot2* and *ggalluvial* (v. 0.12.5). The code for the application is available at https://github.com/meyer-lab-cshl/batman_shiny.

### Amino acid hydrophobicity calculation

To calculate AA hydrophobicity, we used the Wimley-White interfacial hydrophobicity scale [34], which is known to best describe TCR-pMHC AA binding characteristics [58]. This is likely due to the fact that the binding sites of unligated TCRs and pMHC are solvated, and hydrophobic residues at protein-protein interfaces often constitute binding hotspots due to the hydrophobic effect [36, 68]. For AA residues histidine, glutamic acid, and aspartic acid, the above scale provides hydrophobicity of both neutral and charged residues, where we have used the latter. The resulting AA hydrophobicty ordering can be found, e.g., in the horizontal axis of Figure 1F.

### MHC binding predictions

For all MHC-peptide pairs in BATCAVE, we predicted MHC binding using the command-line interface of netMHCpan4.0 [21]. We restricted binding predictions to the full length of the peptide sequence and classified binding based on ‘Rank_BA’ into ‘no’ [> 2% rank], ‘weak’ [0.5-2% rank], or ‘strong’ [< 0.5% rank] binders.

### Training and validation of BATMAN

#### Bayesian hierarchical classifier for TCR activation

We first describe how BATMAN works for a given TCR in within-TCR mode. For classification tasks, BATMAN (Figure 2B) performs Bayesian logistic regression to predict the ordered categorical activation level for the given TCR and peptide, *a* (peptide) ∈ {*a*_*no*_, *a*_*weak*_, *a*_*strong*_}, using the peptide-to-index distance *d* (peptide, index) corresponding to the index peptide of the TCR, using this link function:

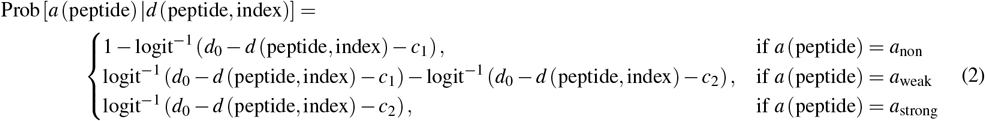

where the inverse logit function is defined as 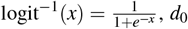 is a constant intercept and *c*_1_ and *c*_2_ are two constant cutpoints with the constraint *c*_1_ *< c*_2_, with the following hyperprior distributions:

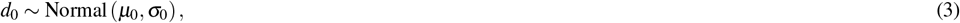

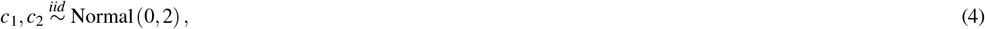

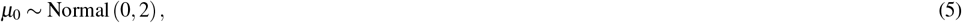

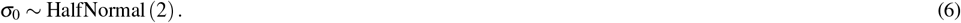

For any peptide-index sequence pair, the peptide-to-index distance *d* (peptide, index) is computed based on position-dependent weights *w* (position) and a 20×20 AA substitution distance matrix *M*:

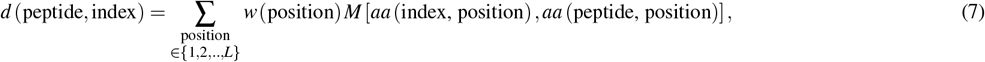

where each element in *M* [*aa* (index, position), *aa* (peptide, position)] corresponds to the substitution of amino acid residue *aa* (index, position) to *aa* (peptide, position) at a given position in the index and peptide sequences, respectively. The diagonal elements of *M* are all zero, such that the distance from the index peptide to itself is zero. BATMAN infers the weights *w* (position) and AA distance matrix elements of *M* [*aa*_1_, *aa*_2_] with *aa*_1_, *aa*_2_ ∈ {A,C,D,…,W,Y}.

Position-dependent weights *w* (position) ∈ [0, 1] with position ∈ {1, 2, .., *L*}, where *L* is the length of the TCR’s index peptide, have the prior:

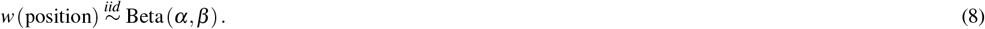

Elements of *M* follow:

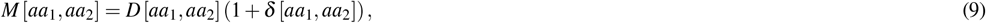

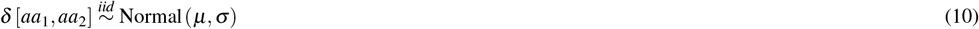

where *D* is a pre-defined AA distance matrix (e.g., BLOSUM100) used for constructing the prior for the inferred AA matrix *M*. The hyperparameters of *d* (peptide, index) have the following, weakly informative hyperprior distributions:

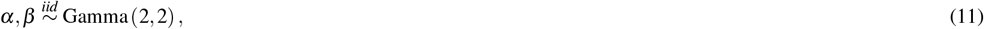

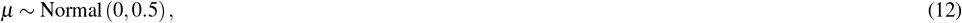

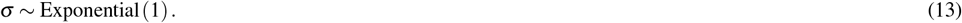

BATMAN is also designed to optionally use pMHC binding information. In such cases, Equation (7) is modified as following:

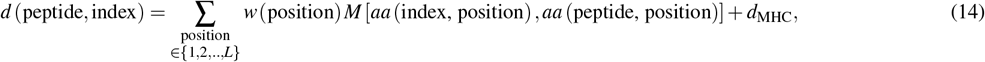

with the scalar *d*_MHC_ assuming one of 3 possible values based on NetMHCPan-predicted pMHC binding level,

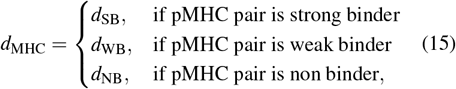

with the the prior distribution

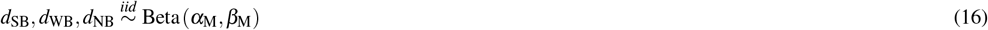

with the hyperparameter prior being

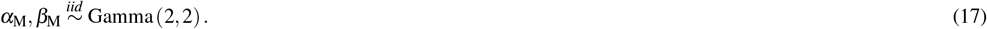

We verified via prior predictive sampling that these assumptions can yield all anticipated outcomes i.e. activation levels.

#### Regression tasks with BATMAN

To use BATMAN for regression tasks of predicting continuous-valued normalized TCR activation *a* (peptide) ∈ [0, 1], we modified Equation (2) to

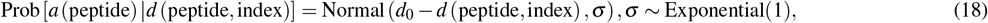

with all other steps being identical as described above for classification tasks. The performance of such an application is shown in Extended Data Fig 5A.

#### Pooling across TCRs for training BATMAN

The hierarchical Bayesian inference set-up allows BATMAN to integrate datasets from multiple TCRs having the same index peptide length (‘pooling across TCRs’). In such cases, the positional weight profiles *w* (position, TCR) and the intercepts *d*_0_ (TCR) are TCR-specific, but have the same prior distributions as specified above, i.e.,

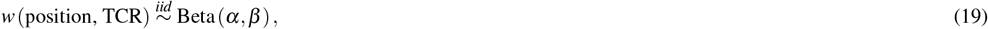

and

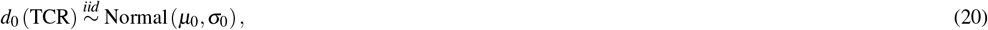

with the hyperparameters *α, β, µ*_0_ and *σ*_0_ having hyperpriors as above. These TCR-specific weight profiles are used to calculate TCR-specific peptide-to-index distances *d* (peptide,index,TCR) similarly as above,

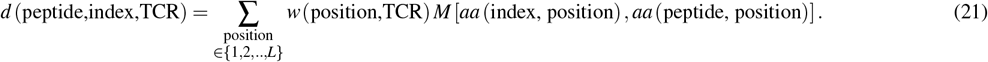

When we used pMHC binding information, we used the following TCR-specific version of Equation (14),

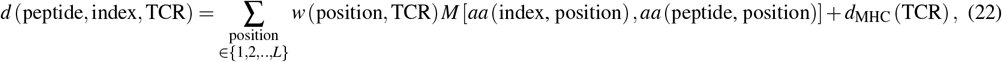

with the TCR-specific quantities *d*_MHC_ (TCR) having the same priors as *d*_MHC_ before.

TCR-specific peptide-to-index distances are consequently used, similar to Equation (2), to construct TCR-specific activation probabilities *a* (peptide,TCR),

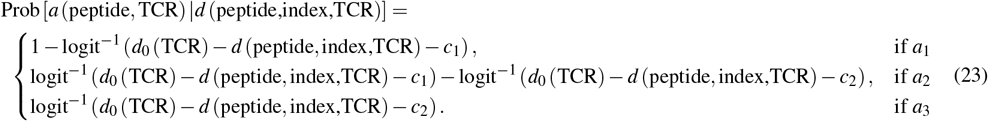

where

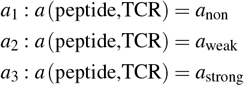

In both within-TCR and cross-TCR cases, pooling was performed over different positions in the peptide sequence, and different elements of the matrix *M*, corresponding to different AA substitutions. Pooling across AA substitutions allowed us to assign *M* [*aa*_1_, *aa*_2_] = *D* [*aa*_1_, *aa*_2_] (1 + *µ*) for AA substitutions absent in the training set but present in the test set.

#### Unpooled BATMAN training

We tested different parameter inference and pooling schemes for BATMAN. For all MHCII TCRs, Figure 2E ‘BATMAN unpooled’ results, all Figure 5A,C (except ‘Pan-TCR’ results) TCRs, Extended Data Figs 5 and 6, and ‘Full_unpooled’ results in Extended Data Fig 7, we did not pool across TCRs, i.e., BATMAN was trained individually for each TCR separately. In these cases, the unpooled inferred weights had a Beta(2, 2) distribution as the prior. We first used conventional AA substitution distance matrices, and performed 5-fold cross validation separately for each TCR, e.g., for 9-mer-binding TCRs using about 144 random peptides from the set of single-AA mutants for the TCR for training, and the remaining 36 for testing.

#### Pooling schemes for BATMAN

In Figure 2D ‘BATMAN (BLOSUM100)’ results, Extended Data Figs 5 and 6, we did not infer the AA matrix, i.e., *M* was set to the indicated AA distance matrix. In other cases where we inferred the matrix *M*, the pre-defined matrix *D* was always chosen to be BLOSUM100, since it performed the best in classification tasks among all the conventional AA distance functions in unpooled training (Extended Data Fig 5B,Extended Data Fig 6). For Extended Data Fig 7 *Symmetric_\** results, we constrained *M* to be symmetric, which showed slightly worse performance than when we did not impose this constraint. Thus, in all other cases, we inferred the full AA matrix. Indeed, the asymmetric part of the inferred full AA matrix was prominent for hydrophobic AA residues (Figure 3E). For plotting the inferred AA matrix in Figure 3E, we divided the inferred matrix by the corresponding values of 1 + *µ*, to make the ratios of inferred matrix elements to BLOSUM100 matrix elements more interpretable.

When pooling across TCRs, for Figure 2D and Figure 2E ‘BATMAN pooled’ results, we pooled across the selected 14 9-mer-binding TCRs and 4 10-mer-binding human TCRs separately. In Figure 2F, Figure 3B, and Extended Data Fig 7 *\*_within* results, we pooled within TCRs specific for an index peptide. In Figure 2D, and Figure 2E ‘BATMAN pooled’ results, BATMAN is pooled across TCRs specific for all index peptides of same length in the benchmarking TCR set of Figure 2A, and across all TCRs specific for all index peptides of same length in Extended Data Fig 7 *\*_across*. BATMAN performance improved by pooling the training data across TCRs (Figure 2E, Extended Data Fig 7), even when inferring TCR-specific weights.

We used pMHC binding information in BATMAN prediction for the results in Figure 2D ‘BATMAN’, Figure 2E,F, and Figure 3. We did not use pMHC binding information in BATMAN-AL, since LOO-TCR performance using pMHC binding had a broader range across TCRs with using pMHC binding (Figure 2D). For multi-AA mutant results in Figure 5A-B, since the experimentally tested multi-AA mutants are biased towards MHC-binders, we did not use pMHC binding information input.

Finally, while in most cases we inferred TCR-specific positional weight profiles, for leave-one-TCR-out tasks (e.g., Figure 2D,F, Figure 4A,B, Extended Data Fig 9 etc) and ‘Pan-TCR’ results of Figure 3A and Figure 5C, we inferred a common weight profile for all TCRs in the training set.

#### Training schemes for BATMAN

For within-TCR validation tasks, we performed 5-fold cross-validation of BATMAN. The folds were stratified by TCR activation levels for classification tasks and TCR activation deciles for regression tasks, and kept identical among all methods (averaged over folds) for comparison.

For TCRs with a sufficient number of peptide examples (≥ 5) of all 3 activation levels to perform 5-fold cross validation, BATMAN classification performance was quantified in terms of 3 pairwise AUCs based on the peptide-to-index distance *d* (peptide, index) of each mutant peptide, calculated using TCR-specific or cross-TCR positional weight profile and AA distance matrix inferred by BATMAN. In such cases (e.g., Figure 2D-F,Figure 4A,B, Extended Data Fig 5 etc) an average of the 3 AUCs are plotted, whereas Extended Data Figs 6 and 7 show individual AUCs. For some TCRs, and test folds, only 2 activation classes were available, where the single AUCs corresponding to the only available activation class pair are plotted. For LOO-TCR tests in Figure 2F restricted among SIINFEKL-specific TCRs, we only used educated TCRs (OT1 and Ed TCRs in Extended Data Fig 2) and discarded low-avidity naive TCRs, for which *pTEAM* performed worse [17].

#### AA distance matrices in prior distribution

To convert conventional AA substitution matrices (*D*^′^ set to BLOSUM_*, PAM_*, Dayhoff, or Gonnet) into distance matrices *D* suitable to be used in priors for BATMAN, we performed the transformation

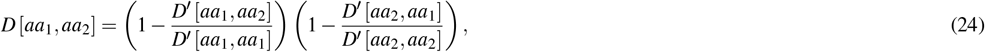

so that the AA distance matrix *D* was always symmetric, with diagonal elements equal to zero. The Hamming matrix had all off-diagonal elements equal to 1.

#### Parameter inference from posterior sampling for BATMAN

BATMAN is implemented in python (v. 3.11.5), using *pymc* (v. 5.10.1) packages. We sampled inferred parameters from approximated posteriors using the “Automatic Differentiation Variational Inference” (ADVI) method [69], with the convergence criterion being that the evidence lower bound (ELBO) loss function did not change by more than 0.1% if the number of iterations was doubled. For unpooled BATMAN training on limited subsamples in Figure 2E, we found that we required a minimum of 30% of within-TCR training fold data for the convergence of estimated BATMAN parameters.

### Other TCR-pMHC interaction prediction methods

#### Training dataset summary of different TCR-pMHC methods

We compared our benchmarking dataset with the training datasets of existing TCR-pMHC interaction prediction methods (Figure 1A). We estimated (1) the total number of TCRs and pMHCs considered by each, and (2) the statistics of all experimentally validated examples of TCR-pMHC interactions spanning their respective full training datasets (Figure 1A). We discarded any subsampling and artificial generation of training dataset (e.g., by random pairing of pMHCs and TCRs, commonly used to generate artificial negative examples). Further method-specific notes on acquisition of training dataset statistics can be found in the Supplementary Notes.

#### Implementation of different TCR-pMHC methods

We tested a subset of pre-trained TCR-pMHC methods on our database. The selection was based on availability of webservers, pretrained models, and ease of installing and running models locally. We trained *pTEAM* in both within-TCR and leave-one-TCR-out modes. For the rest of the methods, we used available pre-trained models on our dataset. Each tested method yielded a continuous-valued TCR-pMHC interaction score for each mutant-TCR pair, which was used to calculate 3 AUCs for classification tasks that were subsequently averaged in the final results. The Supplementary Notes section contains links and summaries of different methods tested, and more technical details on their applications on our database.

#### Implementing pTEAM

A recent method, *pTEAM*, was specifically developed to predict TCR activation by mutants. We implemented *pTEAM* following the description in its source preprint [17]. Briefly, we used Atchley embeddings for index and mutant peptides, and, for leave-one-TCR-out tasks, aligned TCR sequences. These embeddings were used as inputs to random forests with 250 trees for classification and regression tasks, with same folds as BATMAN. Each pairwise AUC was calculated by averaging over two AUCs corresponding to 3 activation level probabilities output from the random forests. We used *R* to align TCR sequences with the *muscle* (v 3.40.0) package and implement the random forests with the *randomForest* (v 4.7-1.1) package. AUCs were calculated using the *multiclass*.*roc* function from the *pROC* (v. 1.18.4) package in *R*.

### Active Learning

#### Active Learning with BATMAN

To initialize parameters, we first trained BATMAN-AL in the leave-one-TCR-out mode, and learned pan-TCR positional weights (*w*_pan-TCR_) and the AA substitution distance matrix (*M*). When applying active learning to a novel TCR, we only modify the TCR-specific positional weights. For all candidate peptides, we used Eqn. (7) to calculate peptide-to-index distances.

Our strategy to select peptides for training was based on a combination of two criteria: (a) picking peptides near decision boundaries so that classifiers can finely delineate classes, and (b) picking diverse peptides so as to cover different regions of peptide space. For (a), if all peptide candidates (single-AA mutants) were sorted based on their peptide-to-index distance, then assuming balanced data, we reasoned that class boundaries would fall at roughly the median distance (for 2-class data) or the two tertile distances (1/3, 2/3 for 3-class data). For (b), we explored diversity in the positions of the selected peptides, as well as diversity in the distances themselves.

Using these intuitions, our BATMAN-AL strategy was as follows. For each position, we first selected one mutant peptide. For roughly half (5 out of 9) the positions — those with the smallest pan-TCR positions weights — we selected the peptide having the peptide-to-index distance closest to the median peptide-to-index distance. For the remaining (4 out of 9) positions, we selected the peptide closest to the first tertile of the peptide-to-index distances (Extended Data Fig 9A). To avoid sampling redundant peptides, when selecting more than one peptide per mutation position, we selected the additional peptides randomly.

To infer positional weights for an unseen TCR, we trained BATMAN in within-TCR mode with the sampled peptides and the index peptide as before, except now the priors for the positional weights (*w*_1_, the positional weight vector for the 1st AL round) are informed by the pan-TCR weights as follows,

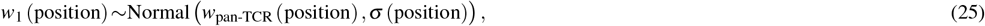

with

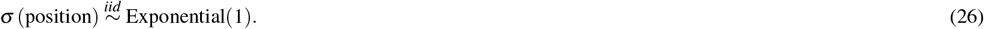

For the next AL round, we sample from the remaining peptides, calculating peptide-to-index distances using the pan-TCR AA matrix and the learned positional weights from the last AL round as a prior:

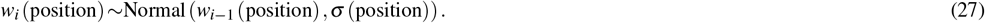

After each AL round, we normalized the learned weights to [0,1], and calculated AUC for the remaining unsampled peptides.

As mentioned above, when selecting more than one peptide per position, our strategy selects the additional peptides randomly. We compared the performance of 4 strategies for selecting random peptides:

1. Peptides are not selected randomly; i.e., peptides are chosen according to their distance from the median or the first tertile of the peptide-to-index distances, depending on the positional weights (i.e., the AL strategy introduced above).
2. All peptides are chosen randomly in each round, while still sampling one peptide per position (“random learning”)
3. All peptides are chosen randomly in even rounds, and in odd rounds, peptides are chosen as per strategy (1) above.
4. All peptides are chosen randomly after the first round, with peptides chosen according to strategy (1) in the first round

For benchmarking performance of AL strategies, among our benchmarking TCR set of Figure 2A, we selected 9-mer-binding TCRs with all positions mutagenized. In all cases involving random peptide selection, we performed AL for 100 realizations of random peptide choice and parameter inference, and averaged the results. The results (Extended Data Fig 9B) showed that out of the 4 strategies, the 3rd strategy exhibited a monotonic increase in median AUC with the number of peptides sampled, and the smallest variance of AUCs, across different TCRs (Extended Data Fig 9B). Thus, we plotted the results corresponding to this strategy in the main text (Figure 4A,B). These results also showed the importance of random peptide choice in AL algorithms. For example, AL without any random peptide sampling did not exhibit monotonic performance increase with more experimental rounds (Extended Data Fig 9B).

### Assessing the relationship between inferred peptide weights with TCR-pMHC structure

For 6 TCRs in the BATCAVE dataset with full peptide mutational scan, there were corresponding PDB entries with the TCRs in complex with either (1) their respective index peptides in the BATCAVE database (TCRs 47BE7, 1E6, and TIL1383I), (2) a single-AA mutant of their index peptide (TCRs NYE-S with BATCAVE index peptide being SLLMWITQ**C** and structure present for SLLMWITQ**V**), (3) the TCR-binding epitope which is a embedded in the longer BATCAVE index peptide (TCRs F5 and F24 with BATCAVE index peptide being FRDYVD**RFYKTLRAEQASQ**E and structure present for RFYKTLRAEQASQ). Since these were the only experimental TCR-pMHC PDB structures available for BATCAVE TCRs, notwithstanding the differences between BATCAVE index peptides and corresponding PDB peptides, we correlated BATMAN positional weights of these TCRs with corresponding structural features with TCR-pMHC interactions.

To compare weight profiles across TCRs, we normalized the positional weights to their TCR-specific maximum. For TCRs F5 and F24, while BATMAN inferred positional weights for all 20 positions of FRDYVD**RFYKTLRAEQASQ**E mutagenized in BATCAVE, PDB structures contained only peptide residues R7-Q19, and thus we restricted our analyses to those for structural analysis.

To correlate the BATMAN inferred positional weights with TCR-pMHC interaction, we analyzed the polar and nonpolar interactions of each peptide residue in the complex with TCR for TCRs with available structural data in the Protein Data Bank (PDB), listed here by TCR name: PDB identifier : NYE-S1:6rpb, NYE-S2: 6rpa, NYE-S3: 6rp9, 47BE7: 7na5, TCR-1E6: 3utt, TIL1383I: 7rk7, TCR-F5: 6cqn, and F24: 6cql. All analyses were performed with UCSF ChimeraX [70] (version 1.8). To identify hydrogen bonding interactions between residues of the peptide to neighboring residues in either TCR, MHC or peptide itself, we used the ‘hbonds’ function which uses atom types and geometric criteria to identify hydrogen bonds. For this, we allowed a distance tolerance of 0.4 Å and an angle tolerance of 20°. To identify all polar and nonpolar interactions involving each of the residues of the peptide to neighboring residues in either TCR, MHC or peptide itself we used the ‘contacts’ function. For this, we allowed a center-center distance of 4 Å. Threshold parameters for interactions represent default cut-offs for the range of the respective interaction [71]. For each pMHC-TCR structure, we then summed the number of polar and non-polar interactions at each peptide position and fit a linear model with interactions as the independent and BATMAN-inferred peptide positional weight as dependent variable.

## Data availability

The fully curated database of TCR-pMHC interactions can be downloaded from https://github.com/meyer-lab-cshl/BATMAN/tree/main/TCR_epitope_database.

## Code availability

Custom analysis code was written in python (version ≥3.10.11) or R (version ≥3.4.0). All analyses and code to re-produce figures in this manuscript are available at: https://github.com/meyer-lab-cshl/BATMAN-paper. The python implementation of BATMAN (‘pyBATMAN’) can be installed from https://pypi.org/project/pybatman/ and run locally. pyBATMAN installation instructions and input file specifications can be found at https://github.com/meyer-lab-cshl/BATMAN/. Example TCR-pMHC input dataset and python script for running pyBATMAN can be found at https://github.com/meyer-lab-cshl/BATMAN/tree/main/run_batman. Interactive Jupyter notebook tutorials on pyBATMAN and BATMAN-AL usage can be downloaded from https://github.com/meyer-lab-cshl/BATMAN/blob/main/run_batman/pyBATMAN_Tutorial.ipynb and https://github.com/meyer-lab-cshl/BATMAN/blob/main/run_batman/BATMAN_AL.ipynb respectively.

**Extended Data Figure 1.**
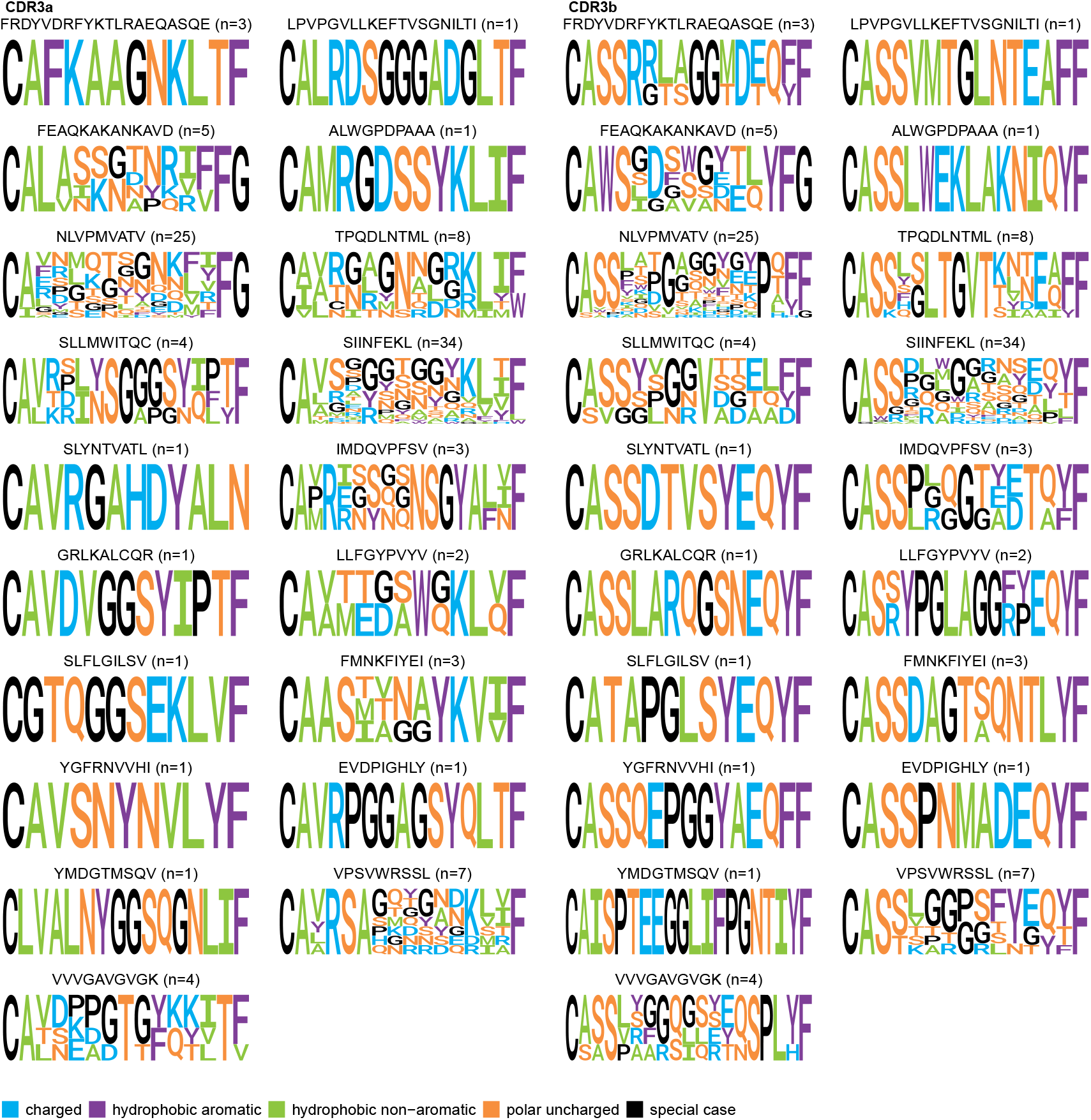
BATCAVE TCR CDR3*α* and CDR3*β* sequence diversity, grouped by index peptides.

**Extended Data Figure 2.**
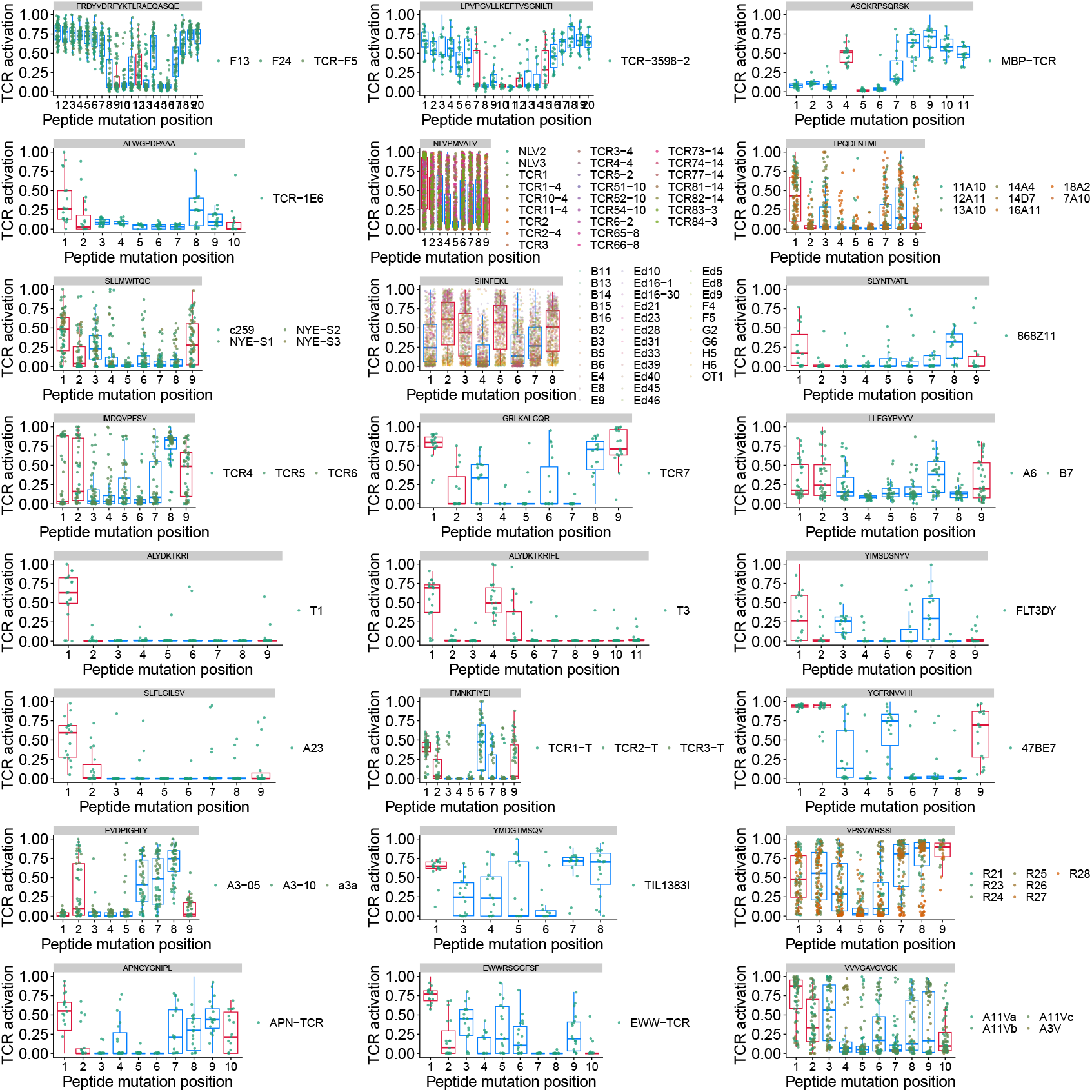
Normalized TCR activation by mutant peptides, grouped by index peptides and mutation position (MHC anchor residues marked red).

**Extended Data Figure 3.**
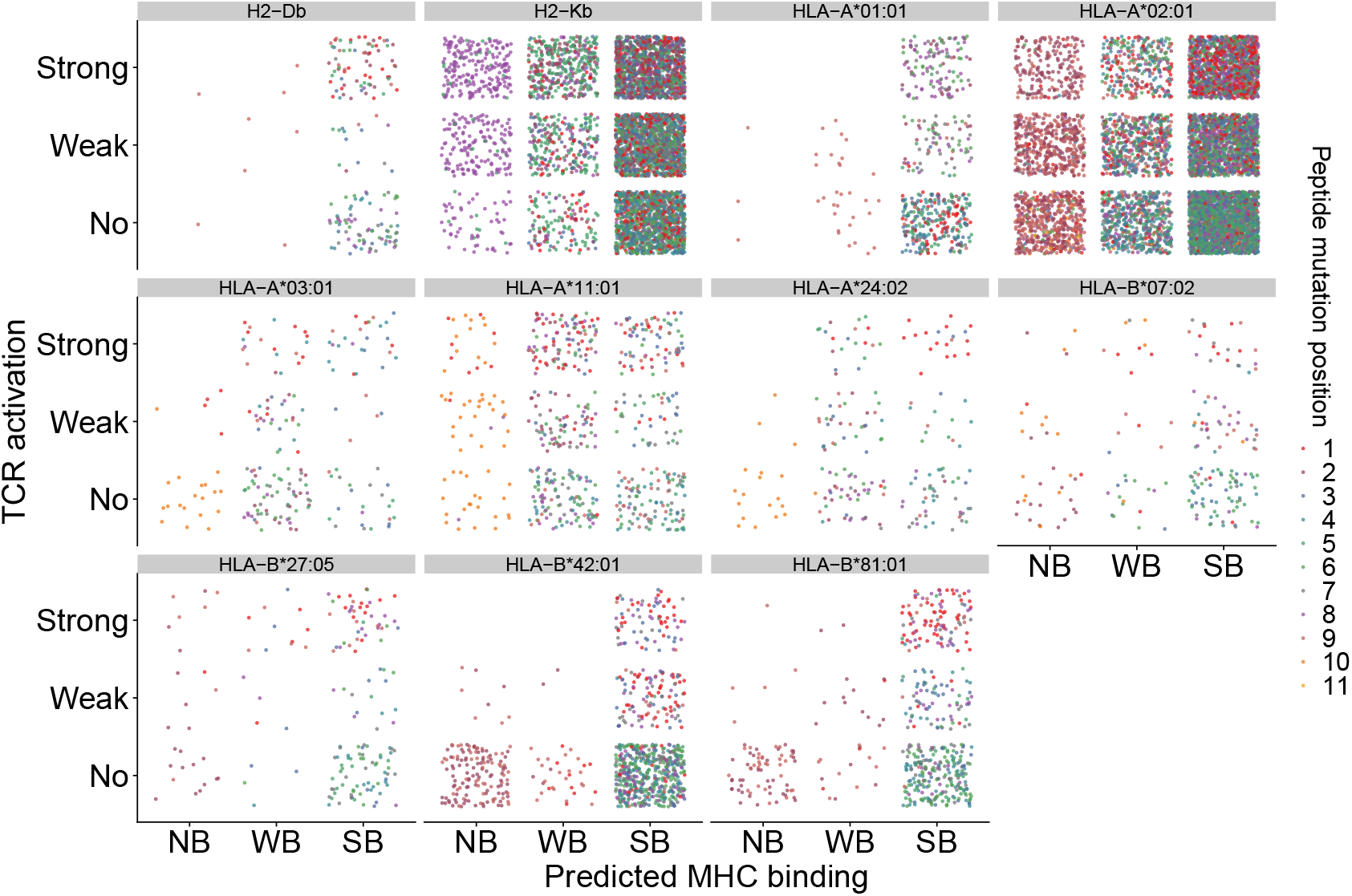
Relationship between pMHC binding and TCR activation in BATCAVE database. Dependency of BATCAVE mutant peptide TCR activation category on NetMHCPan-predicted pMHC binding, grouped by MHCs, for MHCI peptides for which all positions were mutagenized.

**Extended Data Figure 4.**
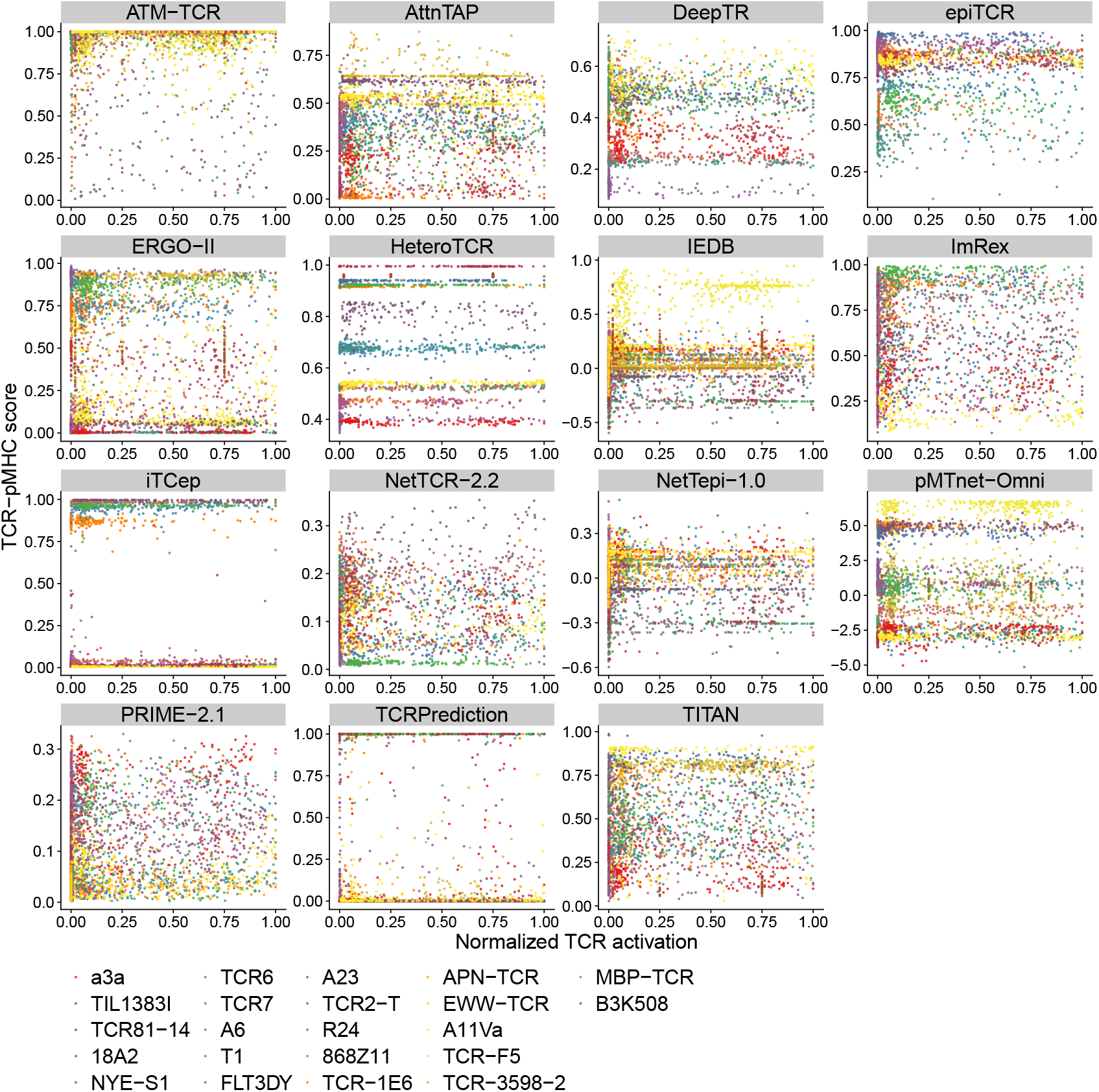
TCR-pMHC scores from different methods do not correlate with TCR activation by mutant peptides. TCR-pMHC interaction scores and normalized TCR activation of mutant peptides for the TCRs selected in Figure 2a.

**Extended Data Figure 5.**
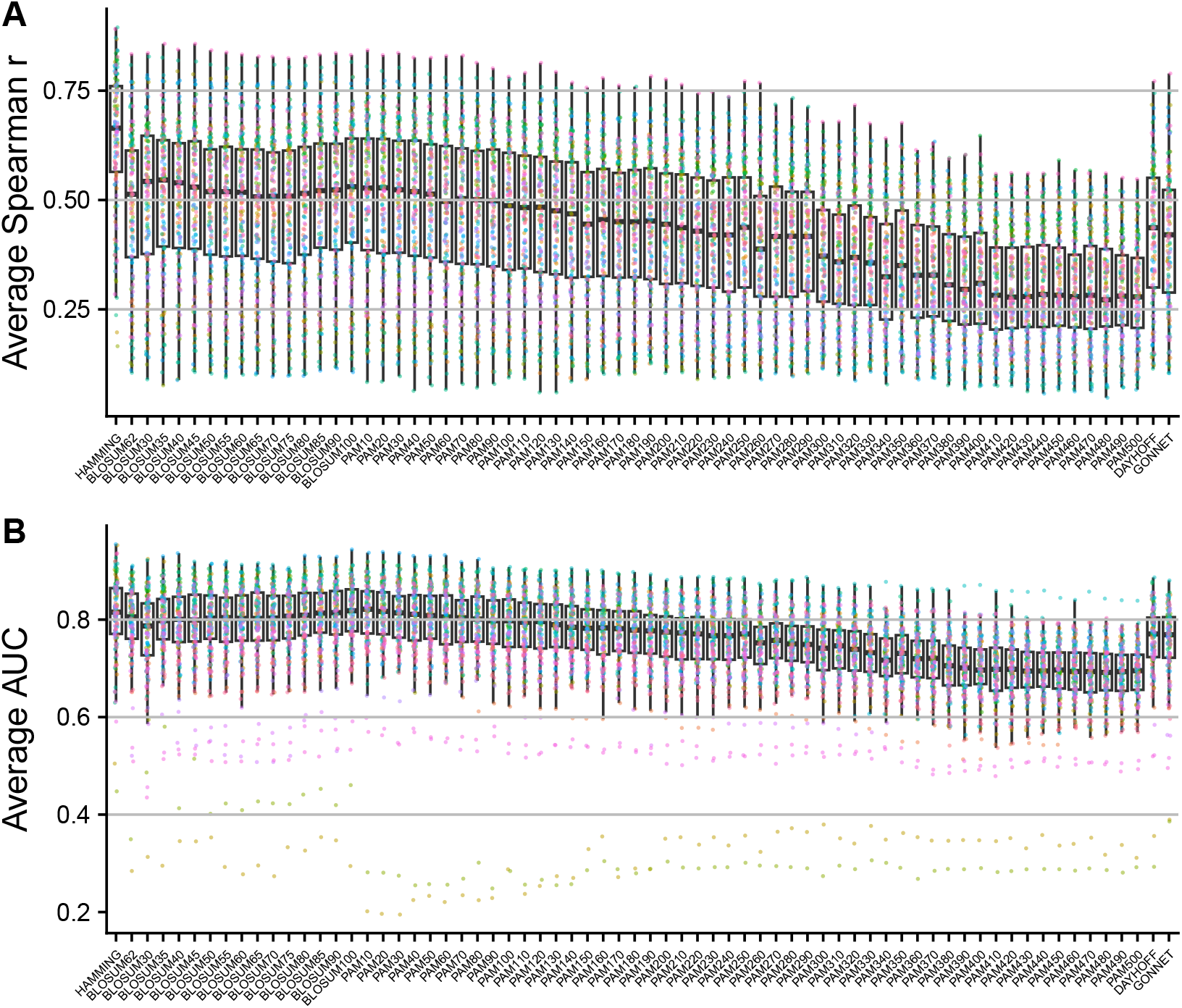
Extended, unpooled performance analyses for BATMAN. **(A)** Classification and **(B)** regression performances in within-TCR tests without cross-TCR pooling using different amino acid distance matrices (points colored by TCRs)

**Extended Data Figure 6.**
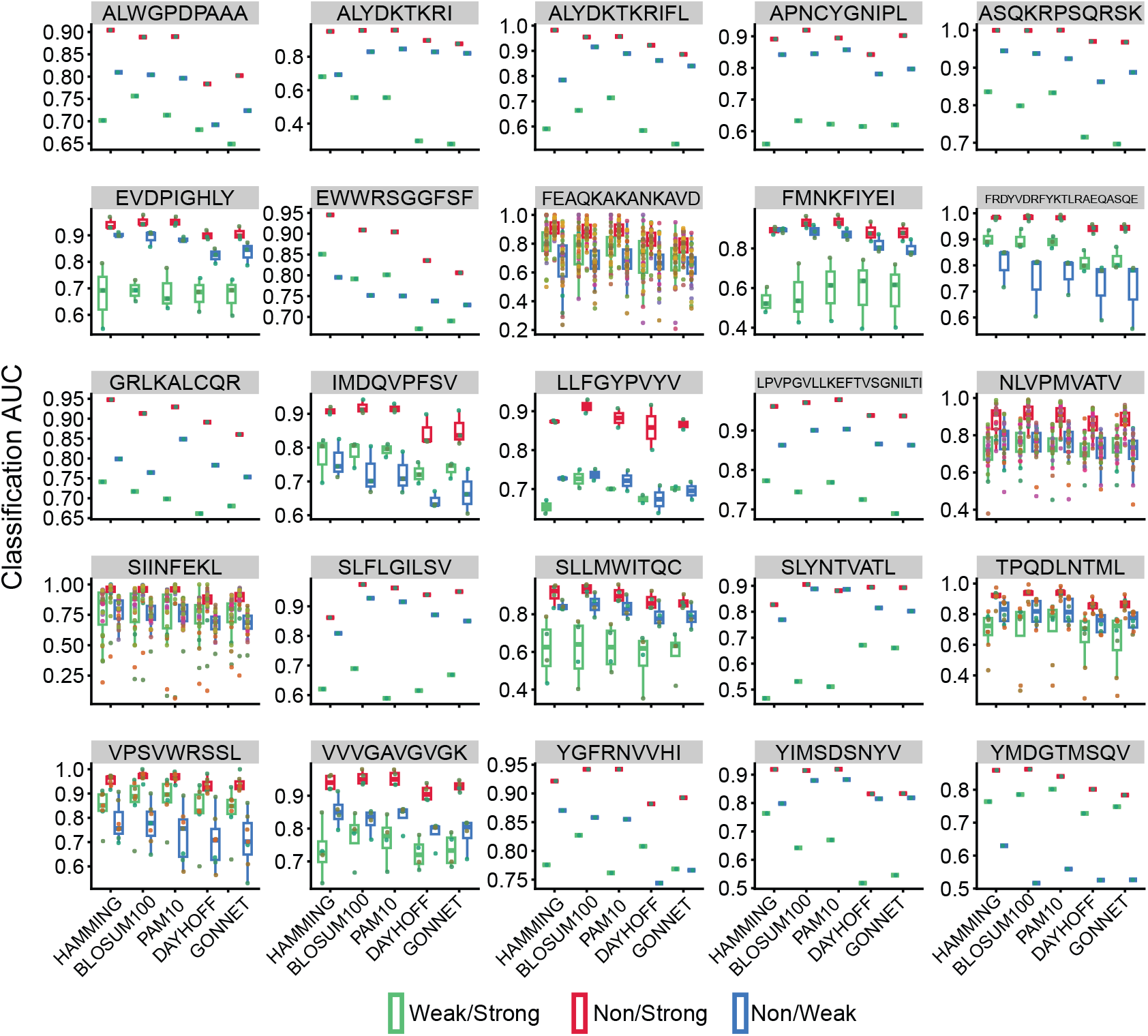
Extended, unpooled performance analyses for BATMAN with selected AA substitution matrices. Pairwise classification AUC for selected amino acid distances for results plotted in Extended Data Fig 5B (points colored by TCRs with the same color scheme as Extended Data Fig 2, grouped by index peptides).

**Extended Data Figure 7.**
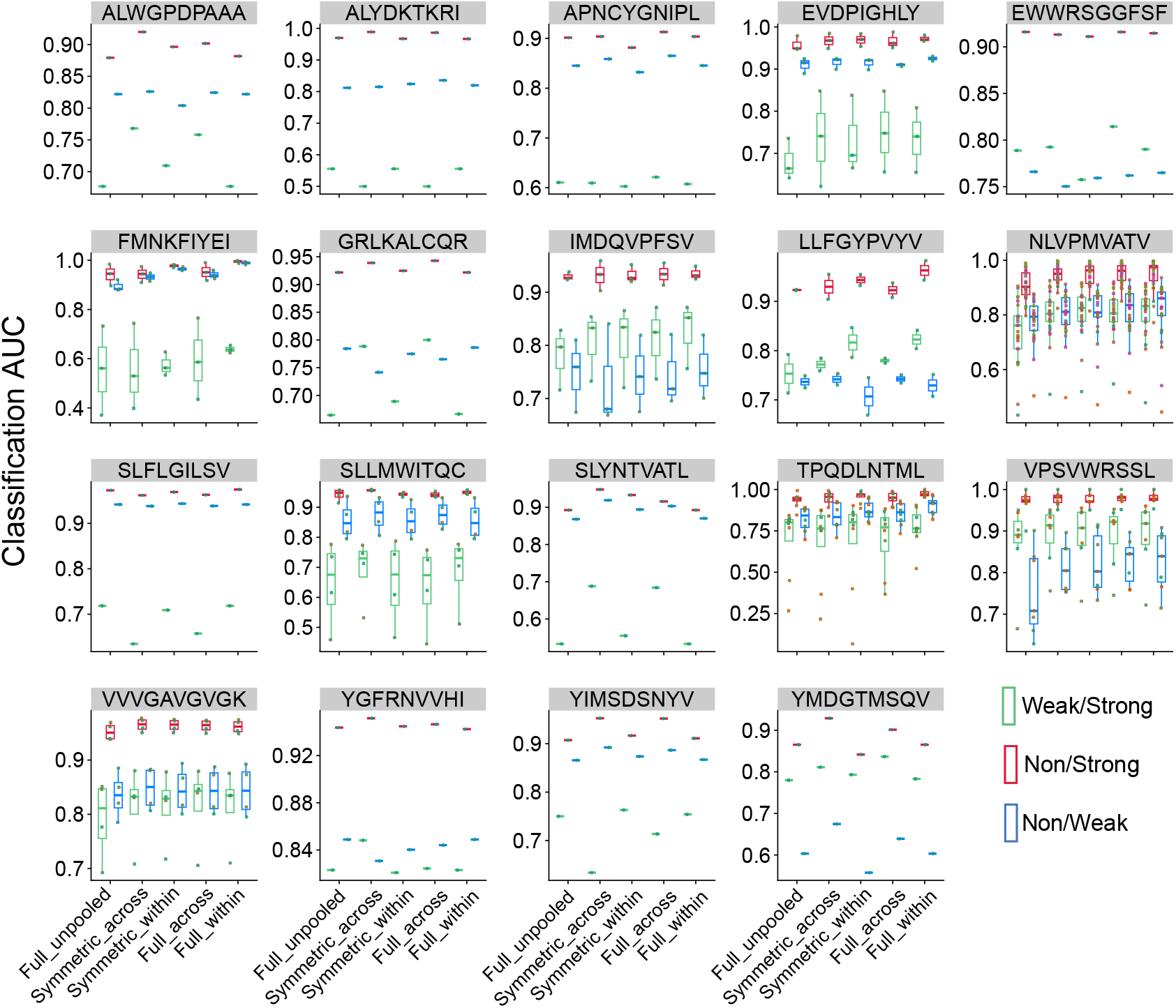
Pooling across TCRs improves within-TCR classification performance. Pairwise classification area under the curve (AUC) for BATCAVE TCRs with 9 and 10 AA long index peptides, with different inferred AA matrices (*Symmetric_\**, and *Full_\**), and pooling modes (*\*_within* TCRs specific for a index peptide and *\*_across* TCRs specific for all index peptides of same length). Unpooled results with inferred Full AA matrices shown for comparison (points colored by TCRs with the same color scheme as Extended Data Fig 2, grouped by index peptides).

**Extended Data Figure 8.**
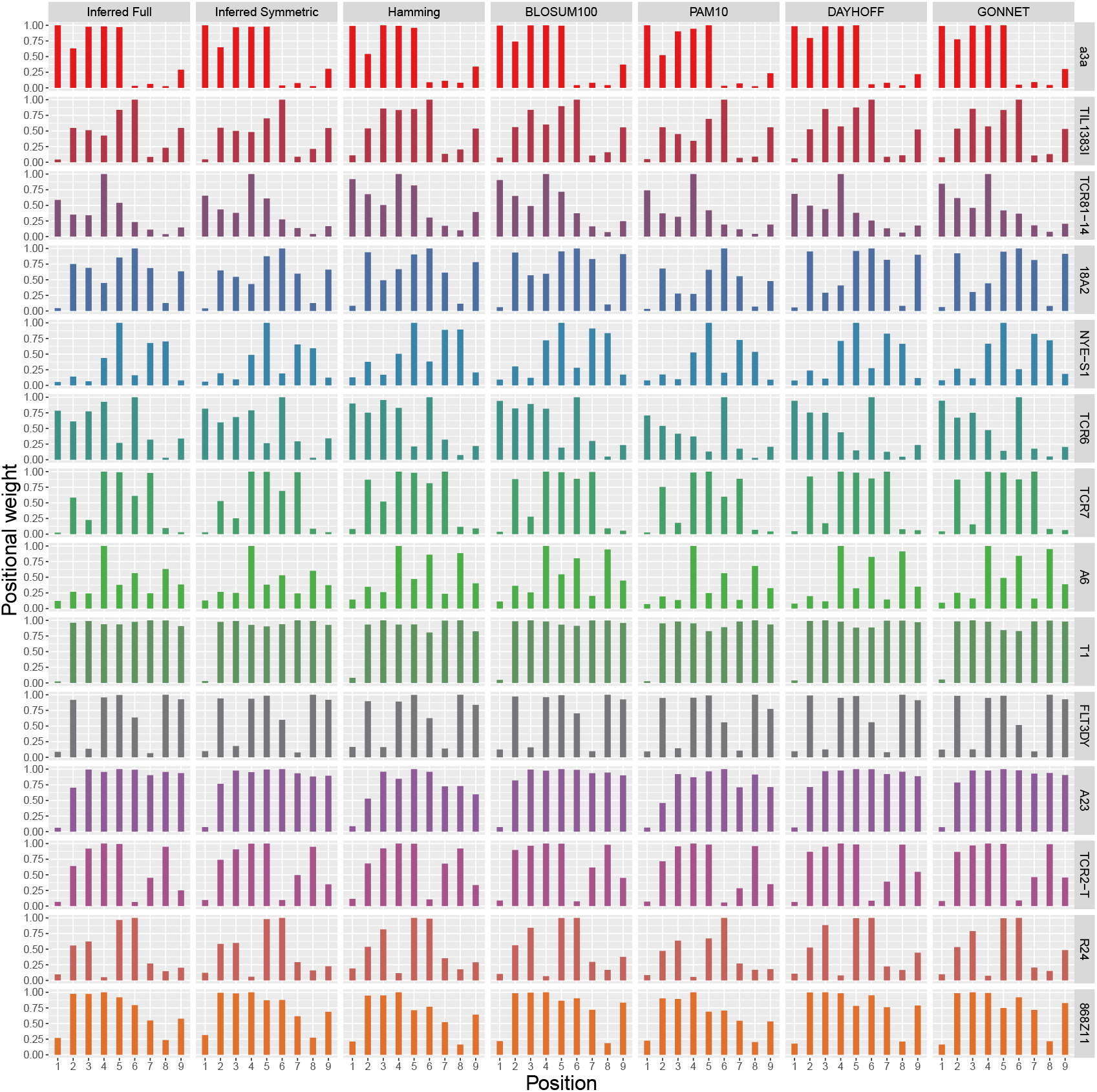
Analyses of inferred positional weights. Positional weights are consistent across different conventional and BATMAN-inferred full and symmetric amino acid distance matrices.

**Extended Data Figure 9.**
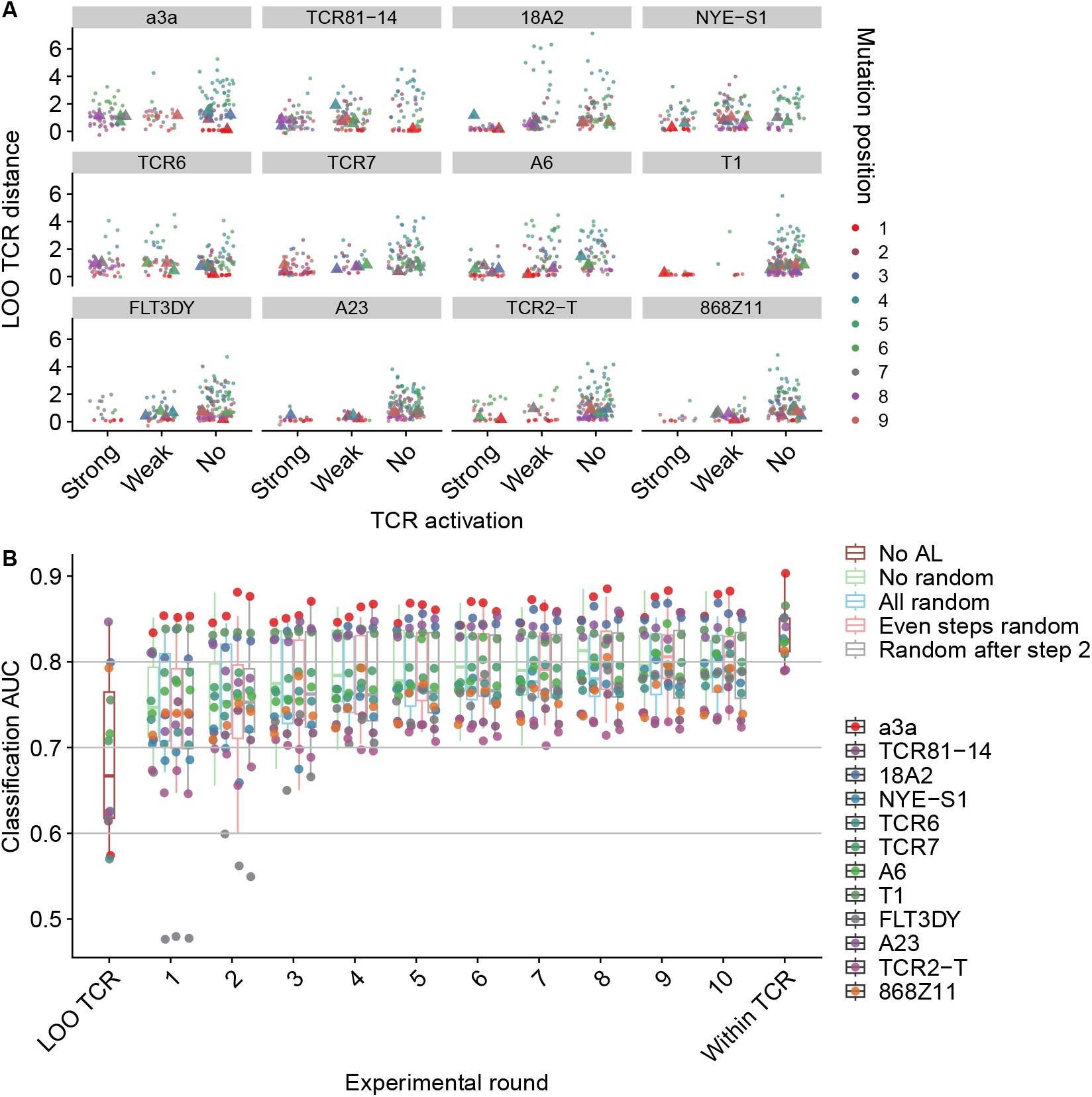
Peptide-to-index distances for mutant peptides are associated with active learning performance boost with BATMAN. **A** BATMAN peptide-to-index distances for single-AA mutant peptides corresponding to individual left out TCRs, using pan-TCR positional weight profile and AA matrix inferred from all other TCRs. The points are separated by TCR activation levels of the peptides, and colored by mutation position. Triangles correspond to the 9 mutant peptides chosen in the first AL round. **B** BATMAN performance for different AL strategies, each sampling 9 peptides per experimental round. For performance comparison, LOO TCR and Within TCR AUCs from Figure 4a are shown.

## Notes

### Competing Interest Statement

The authors have declared no competing interest.

### Summary of Updates

We have significantly extended our database (now covering < 22k peptide pairs) and added many new results and analyses.

