## Supplementary Notes for "Comprehensive epitope mutational scan database enables accurate T cell receptor cross-reactivity prediction"

### Contents

1 Collection of TCR activation and TCR-pMHC binding kinetics datasets 1
2 Training data statistics of TCR-pMHC methods 5
3 Implementation and usage of pMHC-TCR methods 8

### 1 Collection of TCR activation and TCR-pMHC binding kinetics datasets

In this section, we include additional notes on acquisition and processing of TCR activation data from existing publications, grouped by source publication and/or TCR. Code used for acquisition of the data are available on the manuscript github page at: https://github.com/meyer-lab-cshl/BATMAN-paper/tree/main/results_batman/paper_figures/figure_2/2a/tcr_pmhc_methods_predictions.

#### FLT3DY TCR

We sourced the data from Giannakopoulou et al. [1] Figure 1h IFN-$\gamma$ production heatmap. In the mutational scan matrix source data, the recorded IFN-$\gamma$ values corresponding to the unmutated amino acids (denoting the index peptide IFN-$\gamma$ value) are different for different positions on the peptide (likely because they are different replicates of the index peptide). We recorded the average IFN-$\gamma$ value over all occurrences of the index peptide. The FLT3DY TCR sequence is patented.

#### 47BE7 TCR

We sourced the data from Finnigan et al. [2] Fig. 5f IFN-$\gamma$ percentage data, from the “Source Data” spreadsheet associated with the article. Mutant peptide IFN-$\gamma$ measurements have two or three replicates, we take an average over all available replicates. The 47BE7 TCR sequence is recorded from Fig. 2a.

#### a3a TCR

We sourced the mutational scan data for a3a TCR from Kohlgruber et al. [3] Fig. 6b peptide fold enrichment data, from the “Source Data” spreadsheet associated with the article. Note that we skip positions N1, N2, C1, C2 and only consider positions marked 1-9. We also minmax normalized the fold enrichment data to lie between [0,1]. The a3a TCR sequence is recorded from Supplemental Fig. 3(a) of [4].

Additionally, we collected multi-AA mutant peptide activation data for the a3a TCR, with activation labels marked by the source publication or assigned by us as follows:

- The supplementary material Tables S2 and S6 of [5]: We recorded the TCR activation labels as ‘Yes’: SB, ‘Yes,low’:WB, and ‘X’:NB.

- Fig. 6D and supplementary material Table S12 list of [6]: Out of all tested peptides, only the ones marked in color in Fig 6D are recorded as SB, the remainder are recorded as NB.
- Fig. 6a of [3]: Only the exact epitope sequences marked in color are recorded as SB. We neglected all other data in the figure, since the authors performed peptide tile scans involving 90-AA-long sequences, and the exact epitopes, if any, were not identified by them for non-enriched tiles.
- Fig. 5C,E of [4]: Among the tested epitopes in Fig. 5C, the ones with $p < 0.0001$ in Fig. 5E are recorded as SB, the remainder as NB. Additionally, Fig. 6E (and the corresponding peptides listed in Supplemental Table 4), for which, according to the paper, "In addition to epitopes derived from MAGE-A6 and titin, TnT-TCRa3a cells responded to additional peptides (i.e., with higher than 10% activity relative to the MAGE-A3168–176 target), which included the recently reported off-target FAT2, as well nine additional peptides (ANR16, CD166, COG4, CSPG2, FAT1, IL7RA, PLD5, RPAB2, and RUSD2). In contrast, only three and two off-target peptides in addition to the highly homologous MAGE-A6168–176 peptide activated TnT-TCRA3-05 and TnT-TCRA3-10 cells, respectively" So, peptides with <10% activity are recorded as NB, while >10% is recorded as SB.

Note that the peptide EVDPIRHYY had conflicting labels among the above sources. in [6], it is negative (within DMSO control), while in [5] its response is recorded "Yes, low" and it is marked as enriched in [3]. We record it as a WB.

#### APN- and EWW-TCR

We recorded the log2FC data corresponding to Fig 1b,c of Bentzen et al. [7]. The raw data is not available, neither are the heatmaps in vector graphics format. So we saved the heatmaps as webp files, convert them to jpg files, read pixel-wise RGB values using custom Python code and then mapped the RGB values to log2FC values using the colorbar. Since Fig. 1a,b shows that unspecific TCRs have logFC at zero, we set all negative logFC values as 0, and set that as minimum of logFC. We then within-TCR minmax normalized the log2FC values to lie between [0,1]. The publication notes "TCR sequences and expression vectors must be obtained through a material transfer agreement." So we did not include the TCR sequences in our database.

#### KRAS TCRs

We sourced the mutational scan data from NFAT activation heatmaps of Bear et al. [8], found in Fig. 2c. The unnormalized raw data is found in the supplementary "View Supporting data values" Figure 2C sheet. Note that cysteine (C) substitution results are separately recorded in the right of the data sheet. Furthermore, for non-C substitutions, while the alphabetic AA order is maintained for the substituted amino acid, in recording the data in the 18X10 (as opposed to 20X10, or 19X10, excluding only C) grid for each TCR, C and the WT amino acids are missing for each position. So, the set of substituted amino acids is different for each position, and that is not accurately recorded in the grid row amino acid name, since the amino acids cannot be common for all columns. We recorded the data assuming that the amino acid alphabetic order is maintained for each column corresponding to a specific peptide position, with cysteine and the WT AA at that position missing.

Following the legend in Fig. 2c, we minmax normalized the results using the controls for each TCR: Minimum (Jurkat Background) and Maximum (PMA/I Avg), noted separately on the right side of the grid in the spreadsheet. The index peptide activity is also found in the same location. Some peptide activities attain a small negative value after this normalization, we set them to zero in our record. TCR sequences are found from Table 1. In Table 1, the epitope for TCR A11Vc is listed as 8-16V, while for the rest of the TCRs it is 7-16V. Since Fig. 2c performs mutational scan over 7-16 for all TCRs, we record 7-16V as the index peptide for all TCRs.

#### T1 and T3 TCRs

We sourced the data from Ali et al. [9] IFN-$\gamma$ production heatmaps in Figure 2g. In the source data for the mutational scan matrix, the recorded IFN-$\gamma$ values corresponding to the unmutated amino acids (denoting the index peptide IFN-$\gamma$ value) are different for different positions on the peptide (likely because they are different replicates of the index peptide). We recorded the average IFN-$\gamma$ value over all occurrences of the index peptide. The T1 and T3 TCR sequences are patented.

#### TCR* TCRs

We sourced the data from Łuksza et al. [10] Figure 3d,f EC50 heatmaps. We recorded the normalized $-\log_{10}$(EC50) fitted values for mutant peptides, found in Source Data Extended Data Fig. 6, where EC50 was estimated by the fit parameter named "$K_a$". Following the description in the main text [10] with "the inferred EC50's were further clipped to the range of 1E-4 $\mu$g/mL to 1E+4 $\mu$g/mL", we applied the same restriction. We first calculated $a=-\log_{10}$(EC50) for all mutant peptides corresponding to a given TCR, then clipped the values (including $\pm$Inf) to [-4,4] and then mapped all recorded a values to [0,1] for a given TCR. The TCR V$\beta$ and V$\alpha$ sequences were sourced from Extended Data Fig. 4. We aligned known VDJ allele sequences to the TCR sequences to infer the alleles and separate the CDR3 sequences.

### TCR*-T TCRs

We sourced the data from Cai et al. [11] Supplementary Figures S1A and S1B IFNg Mutant/WT ratio barplots. As the source data was not available to us, we opened the figure files in Adobe Illustrator and recorded the total height of the bars in pixels (discarding error bars) and converted them into IFNg values, using the index peptide amino acid bars at each position as the standard.

The TCR CDR sequences were taken from a companion article, Luo et al. [12] Figure 1A. We designated the regions corresponding to sequences starting with "CAAS..." or "CASS..." before "G*G" as CDR3 regions and aligned known VDJ allele sequences to the rest of the TCR sequences to infer the alleles.

### A6 and B7 TCRs

We sourced the raw data from Gejman et al. [13] Figure 3E,F heatmaps for minigene fold change, which was kindly provided to us by the authors. We recorded the normalized $\log_2$ of depletion values for mutant peptides. We first recorded the depletion value for all mutant peptides for a given TCR and then normalized $-\log_2$(depletion) values to map all recorded values between [0,1] for a given TCR. For B7, we clipped the depletion values at 1.5 before normalizing, to minimize the effect of one outlying data point at 1.8 on discrete TCR activation levels.

The TCR sequences were sourced from Supplementary Data Table 3. For A6, we recorded the TCR V$\beta$ and V$\alpha$ sequences corresponding to the bacterial expression construct (for which, $\alpha$ and $\beta$ chain sequences are separately given). We aligned known VDJ allele sequences to the TCR V$\beta$ and V$\alpha$ sequences to infer the alleles and separate CDR3 sequences.

### NYE-S* TCRs

We sourced the raw data from Coles et al. [14] Figures 4D-F IFN$\gamma$ release heatmaps, which was kindly provided to us by the authors. TCR sequences are in Table II of the paper.

### pTEAM TCRs

For ovalbumin-specific ('SIINFEKL') TCRs, we sourced the raw data from Straub et al. [15] Figures 5D and S5B NFAT expression heatmaps, which was kindly provided to us by the authors. The TCR sequences were sourced from another preprint, Dorigatti et al. [16], Supplementary Data 1 and 2. We sourced the NLV-specific TCR data, from Figs. 5A, and S17a NFAT data of Dorigatti et al. [16]. NFAT data and TCR sequence data were sourced from `https://github.com/SchubertLab/TcrPrediction_MutatedAPLs/tree/master/data`. We recorded the unnormalized mutational scan data.

### 868-Z11 TCR

We sourced the TCR activation and TCR-pMHC dissociation constant data from Moritz et al. [17] Figure 6b sheet of the Supplementary Data Sheet. This work reports both binding constant $K_d$ and TCR activation (NFAT-luc2 luminescence) simultaneously for all mutant peptides, of which we record both. We recorded TCR activation value at TCR concentration 100pM, which is the activation threshold for most mutant peptides based on Fig. 6E of the source publication [17]. Note that the amino acid cysteine is missing from the mutational scan due to its tendency to dimerize.

The TCR used here is a single chain TCR, with its construct described in supplementary material S3 of the main publication [17]. We referred to Aggen et al. [18] Supplementary Figure 1 for the full TCR sequence. We aligned known VDJ allele sequences to the TCR V$\beta$ and V$\alpha$ sequences to infer the alleles and separate CDR3 sequences. We designated the sequence "CASSDTVSYEQYF" before the sequence "G*G..." as CDR3$\beta$ in V$\beta$.

Additionally, we sourced multi-AA mutant peptide TCR activation from Supplementary Data "Figure 6d" sheet is taken from [17]. Similar to single-AA mutants, we recorded TCR activation value at TCR concentration 100pM. We subsequently normalized the multi-AA mutant TCR activation values by the maximum of single-AA mutant peptide TCR activation, and assigned activation categories based on the normalized TCR activation following the same convention as single-AA mutants (Figure 1C).

### 1E6 TCR

We sourced the data from Bulek et al. [19] Figure 2 mutational scan heatmap for TNF secretion. The TCR sequence was sourced from the Proten Data Bank entry 5c07, website: `https://www.rcsb.org/structure/5c07`.

### A23 and 1G4 (c259) TCRs

We sourced the data from Foldvari et al. [20] Figures 2A,D IFN-$\gamma$ heatmap (presented in Supplementary Tables 1 and 3). Note that some amino acids are missing in mutational scan of A23 (e.g., Y at position 1). To record IFN-$\gamma$ values, we converted the original PDF file to excel spreadsheet with Adobe PDF exporter and changed the commas (",") to decimal points (".") in the data. We referred to Supplementary Figure 1 of the publication [20] for TCR sequences. We aligned known VDJ allele sequences to the TCR V$\beta$ and V$\alpha$ sequences to infer the alleles and separate CDR3 sequences. Note that the CDR3a region of

1G4 in supplementary reads “CAVRPLYGGSYIP”, which indicates a substitution TS -> LY in the original 1G4 sequence (e.g., found in Fig. 1B of [21], or PDB entry 2BNU, https://www.rcsb.org/structure/2BNU). This variant of the TCR is known as c259 [22], hence we record that name here instead of 1G4.

Additionally, multi-AA mutant peptide activation data for c259 is sourced from Fig. 4B, and Supplementary Data Sheet 1 of [23]. Of these peptides, for the ones in the supplementary, the paper mentions “None of the tested peptides showed evidence of meaningful AKD10R3 cell activation, with a median activation of 1.2% across 65 tested peptides.” So we designate them as NB. For the rest, the paper mentions, “out of 16 peptides with high recognition scores, 44% (7/16) demonstrated at least 75% activation (6, including the original peptide, demonstrated activation above 100%), 27% (4/16) were demonstrated activation in the range of 20–40%, and 33% (5/16) displayed either weak (<15%) activation or no activation at all (Figure 4B).” We designate these three classes as SB, WB, and NB respectively.

While the above single-AA mutation scan data corresponds to SLLMWITQ**C** as the index peptide, for TCR c259, we found matched TCR activation and TCR-pMHC dissociation constant measurements for single-AA mutants of the peptide SLLMWITQ**V** in [24], which we recorded separately for plotting Fig. 1E. For this purpose, we recorded $K_d$ values from Table S2 (Geometric mean) of [24]. The TCR activation data is available from Fig 5C killing assay heatmap of [24]. The raw data is not available, neither are the heatmaps in vector graphics format. So we saved the heatmaps as jpg files, read pixel-wise RGB values using a Python code and then map the RGB values to killing assay values using the colorbar. Cysteine is excluded in the scan at all positions.

#### NLV* TCRs

We sourced the data from Kula et al. [25] mutational scan heatmaps of mutant peptide enrichment in Figure S5. Note that, there were two kinds of enrichment scans done in the paper, with slightly different results: one for 56-aa long fragments (Fig. 5C-D) and one for 9-aa long fragments (Fig. S5), we recorded the latter. Corresponding TCR sequences were recorded from Schub et al. [26] Table II.

#### *A* and *D* TCRs

We sourced the data from Ogunshola et al. [27] mutational scan heatmaps of NFAT luminescence in Figure 5 (raw data was found in Supplementary Data 1 “TCR recognition_normalized" datasheet). TCR sequences were recorded from Figure 4a.

#### TIL1383I TCR

We recorded the mutational scan data for TCR TIL1383I from Fig. 5a ELISA IL-2 data of Singh et al. [28], with the raw data kindly provided by the corresponding author. TIL1383I TCR alpha and beta sequences are recorded from PDB entry 7RK7, Website: https://www.rcsb.org/fasta/entry/7rk7/display. We aligned known VDJ allele sequences to the TCR V$\beta$ and V$\alpha$ sequences to infer the alleles and separate CDR3 sequences.

Additionally, the table in Fig. 6 of [28] contains IL-2 data for multi-AA mutant peptides, with the index peptide IL-2 normalized to 100. So, following our usual normalization scheme (Figure 1C) we record the activation categories based on IL-2 measurements as follows: <10: NB, 10-50: WB, and $\geq$ 50: SB.

Finally, we derived TCR-pMHC dissociation constants $K_d$ for mutant peptides from the $\Delta\Delta G$ heatmap of Fig. 5b. The raw data are not available, neither are the heatmaps in vector graphics format. So we saved the heatmaps as webp converted to jpeg file, read pixel-wise RGB values using a Python code and then map the RGB values to $\Delta\Delta G$ values using the colorbar. From $\Delta\Delta G$ data, we calculated $K_d$ by using the equation $K_d = \exp(\Delta G/RT)$, with $\Delta G = \Delta G_{WT} + \Delta\Delta G$, where $\Delta G$ of WT peptide is -6.25 kcal/mol from Fig. 6 table and T is assumed to be 298K, with $RT = 0.592$ kcal/mol (this also matches Fig. 6 $\Delta G$ and $K_d$ conversions).

#### A3-05 and A3-10 TCRs

We recorded the %CD69 data corresponding to Fig. 6a in the paper Vazquez-Lombardi et al. [4]. The raw data is not available, neither are the heatmaps in vector graphics format. So we save the heatmaps as jpg files, read pixel-wise RGB values using a custom Python code and then map the RGB values to %CD69 values using the colorbar. Note that we skipped a3a mutational scan data, which is already recorded from [3] as described above, and shows very similar results between Fig. 6b of [3] and the corresponding values in Fig. 6a of [4]. The relevant TCR sequences are patented.

Additionally, Fig. 6E (and the corresponding peptides listed in Supplemental Table 4) of [4] contain A3-05 and A3-10 TCR activation data for multi-AA mutant peptides. For recording the TCR activation levels of these peptides, we followed the same scheme as a3a TCR multi-AA mutant peptide activation data, sourced from the same figure and described above.

#### 3K-specific TCRs

We sourced the mutational scan data from Figure 2 of Huseby et al. [29]. The data is presented as a categorical heatmap of 3 T cell response classes. We recorded, with index peptide activity set to 1,

- White pixels, >50% of index peptide response, as 0.75, to classify them as SB,
- Yellow pixels, 5-50% of index peptide response, as 0.25, to classify them as WB,
- Red pixels, <5% of index peptide response, as 0.02, to classify them as NB

For a subset of above TCRs, the TCR-pMHC $K_d$ values for mutant peptides are available from [30]. For a subset of these peptides, the $K_d$ values are explicitly denoted in Table 1, which we record directly. For the rest, we extracted the $\Delta\Delta G$ data from the bar heights in Fig. 3 and converted them to $K_d$ values using the formula $K_d = \exp(\Delta G/RT)$, with $\Delta G = \Delta G_{WT} + \Delta\Delta G$ with $\Delta G_{WT}$ recorded from Table 1, and T assumed to be 298K, with $RT$=0.592 kcal/mol.

We sourced the TCR sequences from Supplementary Table 1 of [30] and Supplemental Experimental Procedures of [29].

#### MBP-TCR

We recorded the proliferation data of Grogan et al. [31] Fig. 1a. The index peptides show slightly different values for different replicates, which we averaged over. TCR sequence was not available.

#### TScan-II TCRs

We recorded the mutational scan data from Fig. 2D-F and 4E peptide fold enrichment data of Dezfulian et al. [32], found in Supplementary tables S1 and S2 online with the article. For TCRs F5, F13, and F24 we skipped positions 1, 2, 23, and 24 since the epitope sequence is denoted as FRDYVDRFYKTLRAEQASQE [32]. In the data, peptides sequence replicates are reported as multiple entries, we averaged over all occurrences of each peptide as part of the full 56-aa tile SILDIRQGPKEP**FRDYVDRFYKTLRAEQASQE**VKNWMTETLLVQNANPDCKTILKA. For TCR-3598-2, data, peptides sequence replicates are reported as multiple entries, we average over all occurrences of each peptide as part of the full 56-aa tile AMPFATPMEAELARRSLAQDAPP**LPVPGVLLKEFTVSGNILTI**RLTAADHRQLQLS. We minmax normalized the enrichment data to lie between [0,1] for each TCR separately. TCR sequences for TCRs F5, F13, and F24 are taken from Supplementary table S11 of [33], and that of TCR-3598_2 from Table 1 of [34].

#### Additional publications for future consideration

A number of publications report more mutational scan datasets for TCR activation, but the raw data from these papers were not readily available. We plan to include them in our database when they become available. These include for instance Bentzen et al. [7], Border et al. [35], etc.

### 2 Training data statistics of TCR-pMHC methods

To plot Figure 1a, we collected the number of TCRs and pMHCs used by different TCR-pMHC prediction methods, and the number of experimentally-verified positive and negative examples of TCR-pMHC interactions in all cases. In all cases, we disregarded the (usually negative) data that is artificially generated (e.g., by randomly pairing TCR and pMHC) without experimentally validating TCR-pMHC interaction. Note that the numbers below are approximate since we did not consider subsampling, or splitting data into folds during training. Thus, the number of experimentally validated TCR-pMHC interactions broadly corresponded to aggregated numbers over all databases considered in the respective methods. Code and input training datasets to calculate training dataset statistics of different methods, if applicable, are available at: `https://github.com/meyer-lab-cshl/BATMAN-paper/tree/main/results_batman/paper_figures/figure_1/1a`

#### BATMAN

For our curated database, non-activators (i.e., normalized TCR activation <0.1) were treated as negative instance and the rest were treated as positive instances.

#### pTEAM

The TCR-pMHC data is from mutational scans of SIINFEKL, a human neoantigen, and NLV-peptides in [16]. We used normalized activations of 46.9, 66.09, and 40 are as thresholds for defining binders as per the paper (data source: datasets S1-S4 files). Unique peptides: all 153 SIINFEKL mutants + 134 neoantigen mutants + 172 NLV-mutants. Unique TCRs: (15+21) Ova-TCRs + 7 neoantigen-TCRs + 20 NLV-TCRs.

#### IEDB

Main paper is Calis et al. [36] where the training data is found in 2 Dataset files. Since the dataset does not use any TCR information, TCR count is recorded as 0 and the same convention is followed for all other immunogenicity models below.

#### PRIME 2.0

Main paper is Gfeller et al. [37], where the training data is in supplementary table S4. From the dataset, "random=1" datapoints were discarded as they were not experimentally validated (only about 6000 data points were experimentally validated, corresponding to "random=0").

#### ImRex

Main paper is Moris et al. [38]. We recorded the training statistics based on the quote [38] "This dataset was reduced to 19,842 unique CDR3-epitope pairs by selecting only human TCR sequences". We discarded negative data from unspecific TCR sequences, which led to only positive examples remaining. To estimate representative TCR numbers, we downloaded an example train data from the GitHub link `https://github.com/pmoris/ImRex/blob/master/models/pretrained/2020-07-30_11-30-27_trbmhci-shuffle-padded-b32-lre4-reg001/iteration_no_validation/train.csv`, filtered it for y=1 (corresponding to known binders, whereas y=0 corresponds to data generated by random shuffle) and found the number of unique CDR3s and peptides.

#### pMTnet

Main paper is Lu et al. [39]. We recorded the training statistics based on the quote [39] "We collected a total of 32,607 pairs of TCR–pMHC bindings from a series of peer-reviewed publications... We created ten times more negative pairs by randomly mismatching the TCRs and pMHCs of these 32,607 pairs.", i.e. this dataset does not contain any experimentally verified negative data. We downloaded the training data from the GitHub link `https://github.com/tianshilu/pMTnet/blob/master/data/training_data.csv` and counted 29226 unique CDR3 sequences and 429 unique peptides.

#### epiTCR

Full experimentally validated training data was available for both negative and positive TCR-pMHC pairs. Main paper is Pham et al. [40]. Training dataset was downloaded from the GitHub link `https://github.com/ddiem-ri-4D/epiTCR/blob/main/data/categories/full-training-with-categories.csv.zip` following which we counted the number of binding and non-binding CDR3$\beta$-pMHC pairs.

#### ERGO II

Main paper is Springer et al. [41]. The training dataset does not contain any experimentally validated negative data. Since the model trained on the McPAS dataset performed better than the one trained on the VDJdb dataset in both the original publication and on our own dataset, we recorded the statistics for the McPAS-trained model. Following the dataset reported in the supplementary table of the main paper [41], we recorded the number of unique CDR3b sequences as the number of unique TCRs in the training data: 9490, and number of unique peptides=319, number of positive sample=11630.

#### NetTCR-2.0

Main paper is Montemurro et al. [42]. We recorded the statistics for prediction mode 'paired TCR' which has superior performance in internal mode comparisons [42]. We recorded the training statistics based on the quote [42] "Positive data points were taken from ... 3859 unique binding pairs were identified from IEDB and 2843 from VDJdb. These provided 4598 unique CDR3$\alpha$-/$\beta$-peptide interactions with 276 different peptides specific to allele HLA-A*02:01.

Negatives were derived from 10X. Using the same restrictions as for the positives (CDR3 length between 8 and 18 AAs, peptide length 9, and peptides specific for HLA-A*02:01), 627,323 unique data points with 0 UMI counts to all the tested peptides were identified. These contained 33,017 unique TCRs tested against a set of 19 different peptides. In total, 17 of these overlapped with the peptides in the positive data set."

Thus, total number of TCRs=4598+33017=37615, with 4598 positive and 627,323 negative instances, and 276+19-17= 278 peptides.

#### TCR-BERT

Main preprint is Wu et al. [43]. We disregarded the pre-training and TCR embedding learning data, and only recorded the numbers of LCMV peptide specific TCR sequences (so 1 unique peptide considered). We used the following quote to record training statistics [43]: "Overall, this results in n=17,702 unique TRA/TRB pairs with consistent labels that we use for model training and evaluation. Among these, 13% (n=2306) are observed to have mid or high binding – we consider these "positive" examples of TRA/TRBs binding GP33."

#### TCRdist

Main paper is Dash et al. [44]. No negative data was considered, training data is summarized in Extended Data Table 1. We recorded from the training data summary that Total number of TCRs = total number of positive examples = sum of Number of clones for 10 peptide-specific repertoires $= 117 + 305 + 324 + 642 + 87 + 158 + 291 + 76 + 275 + 61 = 2336$.

#### TITAN

Main paper is Weber et al. [45]. All negative data was generated by shuffling, and so were disregarded in reporting the training statistics. We used the following quote [45] to record training statistics: “With this procedure, we build a training dataset of 46,290 examples, 50% of which are positive, encompassing 192 different epitopes."

#### DeepImmuno

THis is a general immunogenicity prediction method, so no TCR information is involved. Main paper is Li et al. [46], from which we use the following quote to record training data statistics : “Specifically, 8971 data instances were retained in the final dataset, among which 4059 were positive reactive instances and the remaining 4912 were negative".

#### MixTCRpred

All negative data was generated by shuffling, so we disregarded them in calculating training data statistics. To record training data statistics, we used the quote from the Source preprint Croce et al. [47]: “For further analysis, only pMHCs with ten or more binding TCRs were considered, resulting in a total of 17,715 $\alpha\beta$TCRs interacting with 146 pMHCs (Figure 1C and Table S1)". We also downloaded the full training data from the GitHub link `https://github.com/GfellerLab/MixTCRpred/blob/main/full_training_set_146pmhc.csv` and confirmed that the total number of data points is equal to 17,715.

#### PanPep

Main paper is Gao et al. [48]. Negative data was generated computationally, so we disregarded them in recorded training data statistics. For the rest, we used the following quote [48] to record training data statistics: “As a result, the base dataset contains 699 unique peptides and 29,467 unique TCRs with 32,080 related peptide–TCR binding pairs considering the cross-reactivity of peptide–TCR binding."

#### BigMHC

We collected statistics for the “Immunogenicity Training" mode and used the quote “BigMHC transfer learned only on the nonrandom pMHC data, of which 1,580 are positive and 5,293 are negative" from the main paper Albert et al. [49] to record training data statistics.

#### catELMo

Main preprint is Zhang et al. [50]. Negative data is generated randomly, so we disregarded them in calculating the training data statistics. For recording statistics of the positive data, we used the quote [50] “Altogether, we obtained 150,008 unique TCR-epitope pairs known to bind to each other having 140,675 unique TCRs and 982 unique epitopes."

#### ATM-TCR

Main paper is Cai et al. [51]. Negative data is generated randomly, so we disregarded them in calculating the training data statistics. For recording statistics of the positive data, we used the quote [51] “This resulted in the primary dataset containing 128,142 unique TCR-epitope pairs, with 931 unique epitopes and 119,984 unique TCRs."

#### GLIPH

No negative data was considered. The main paper, Glanville et al. [52] quotes: “In total, the training set consisted of 2,068 unique TCRs of known specificity (Supplementary Table 1)". We found that the Supplementary Table 1 (sheet=“Curated") had 1,973 unique CDR3s and 7 unique peptides, with 2,067 CDR3-peptide pairs in total.

#### Repitope

Main paper is Ogishi et al. [53]. Even though they consider both TCR and peptide sequences, they are not matched in the training data. We included the statistics from the training data reported in Supplementary data sheet 2. We neglected immunogenicity label contradictions in data and focused on the immunogenicity column only, counted positives and negatives and added them up.

#### TCRAI

Source paper is Zhang et al. [54]. We downloaded the training data from the GitHub link `https://github.com/regeneron-mpds/TCRAI/blob/main/data/CNN-prediction-with-REGN-pilot-version-2.csv` and recorded the number of unique TCRs, peptides, and TCR-peptide pairs (positive and negative were based on the ‘binds’ column).

#### DeepTCR

Main paper is Sidhom et al. [55]. DeepTCR has a (1) TCR encoding mode, trained on antigen-specific TCR sequences (only positive examples), and (2) a UMI regression mode, trained on the 10X data. For mode (1) (TCR encoding), the publication quotes [55] "we collected data for tetramer-sorted antigen-specific cells for nine murine (Db-F2, Db-M45, Db-NP, Db-PA, Db-PB1, Kb-M38, Kb-SIY, Kb-TRP2, Kb-m139) and seven human (A1-CTELKLSDY, A1-VTEHDTLLY, A2-GILGFVFTL, A2-GLCTLVAML, A2-NLVPMVATV, B7-LPRRSGAAGA, B7-TPRVTGGGAM) antigens where the ground truth label corresponds to a particular antigen specificity for an individual sequence". This set comprises all the positive data they considered. To estimate the number of TCR sequences used, we counted the total numbers found in the GitHub folders https://github.com/sidhomj/DeepTCR/tree/master/Data/Murine_Antigens and https://github.com/sidhomj/DeepTCR/tree/master/Data/Human_Antigens to find $(117+291+305+324+642+158)+(776+655+456+600)+(410+435+347)+87=5,603$ murine and $25+271+(275+588)+(76+739)+(61+166)+214+65=2,480$ human TCRs and an equal number of positive TCR-pMHC binding. For mode (2) (regression), the publication quotes [55] "a second single-cell dataset published by 10x Genomics where the binding to cognate T cells of 57,229 unique $\alpha/\beta$ pairs to 44 specific pMHC multimers and 6 negative controls was characterized". This dataset was downloaded from the GitHub link https://github.com/sidhomj/DeepTCR/tree/master/Data/10x_Data and has continuous count information on 57,228*44 TCR-pMHC pairs. Out of them, we designated negative as count=0 (n= 2,562,701) and positive as count > 0 (n= 298,699). A total of 9 murine and 7+44+6-5 (common) =52 human antigens were considered. In reporting the statistics of the deepTCR model, we combined the number of positives from the two modes (5603+2480+298699=306782). Total number of TCRs were also combined (5603+2480+57228=65311).

#### TEINet

No experimentally validated negative data was used. To record training data statistics, we used the following quote "At last, we constructed a large dataset with 44,682 pairs of TCRs and epitopes, among which 41,610 TCRs are linked to 180 epitopes" from the main paper Jiang et al. [56].

#### SwarmTCR

No experimentally validated negative data was used. To record training data statistics, we used the following quote "In total, our default SC dataset comprised 1447 TCRs (for complete dataset counts see Additional file 1: Table 4)", from the main paper Ehrlich et al. [57]. We found that Table 4 has 83 peptides.

#### TCRGP

No experimentally validated negative data was used. To record training data statistics, we used the following quote, "The VDJdb data and the Dash data have some overlap for TCRs specific to three epitopes: In the VDJdb data 34 (27 unique) of the 413 (242 unique) TCRs for pp65495-503, 30 (27) of 299 (152) for BMLF1280-288, and 74 (61) of 239 (138) for M158-66 can also be found from the Dash data" from the main paper Jokinen et al. [58]. So the data sources are VDJdb and Dash data. We looked into the GitHub folder https://github.com/emmijokinen/TCRGP/tree/master/training_data/paper for data (files names marked with "unique") and found $138+474+120+234+286+542+526+586+1168+174=4248$ unique Dash TCRs (corresponding to "three epitopes from humans and seven epitopes from mice" [58]) and $278+78+120+156+190+276+304+106+92+118+116+208+130+282+396+484+100+298+58+80+240+104+610=4,824$ unique VDJdb TCRs (23 peptides). So there was a total of $4,248+4,824-27-27-61=8,957$ unique TCRs, and $10+23-3=30$ peptides combining two datasets.

### 3 Implementation and usage of pMHC-TCR methods

The project GitHub has input and output TCR datafiles for 15 TCR-pMHC methods tested in Figure 2A.

#### IEDB

For MHCI peptides, we uploaded peptide sequences separately for each MHC to the Webserver https://nextgen-tools.iedb.org/pipeline?tool=tc1 [36] with prediction model "Class I pMHC Immunogenicity" with default (1,2, C terminal) positions to mask, the corresponding MHC allele selected, and Peptide Length(s) set "as is". Prediction for HLA-B81:01 returned errors, so we could not use them. For MHC II data (binding predictions only), we uploaded peptide sequences separately for each MHC to Webserver http://tools.iedb.org/mhcii/ [59] with "Prediction method:" set to "NetMHCIIPan 4.1 EL", the corresponding MHC allele selected, and length set to "as is".

#### PRIME 2.0

We submitted pMHC information to the webserver at http://ec2-18-188-210-66.us-east-2.compute.amazonaws.com:3000/ [37, 60], separately for each MHC allele, and indicated the corresponding MHC allele in the submission form for each submission. We used the output peptide rank ('%Rank_bestAllele' column in the output file) to calculate the AUCs in Figure 2B. The best MHC allele was same as the allele in our database in all cases. No mouse or MHCII predictions were available, only HLAI predictions were available. Among different outputs, we used the PRIME score for predicting TCR activation.

#### pMTnet_Omni

Prediction is available only for TCRs with at least the CDR3$\beta$ sequence. For TCRs with full sequence available, we formatted the data according to https://pmtnet-omni-document.readthedocs.io/en/latest/input_format/index.html (more details in the code to generate input) and used the online webserver of pMTnet_Omni at https://dbai.biohpc.swmed.edu/pmtnet/analysis-omni.php [61] for predictions. We used the logit column as prediction. For the TCR2-T TCR with only the CDR3$\beta$ sequence available, we used pMTnet (v1), and used the webserver https://dbai.biohpc.swmed.edu/pmtnet/analysis-base.php [39].

#### NetTepi

We submitted TCR-pMHC data to the webserver at https://services.healthtech.dtu.dk/services/NetTepi-1.0/ [62]. Only HLAI predictions were available. Among them, no predictions existed for HLA-B81:01. We uploaded peptide sequences separately for each HLA and for 9mers and 10mers to the webserver at https://services.healthtech.dtu.dk/services/NetTepi-1.0/, set default values for Relative weight on stability prediction: 0.16, and Relative weight on T cell propensity prediction: 0.10 and indicated the respective peptide lengths and MHC alleles. T Cell Propensity score gave better AUC than combined score, so we used the former score.

#### NetTCR-2.2

The predictions require TCR CDR1$\alpha$, CDR2$\alpha$, CDR1$\beta$, and CDR2$\beta$ sequences, which we sourced from IMGT database for different TRA and TRB genes in data (e.g., from https://www.imgt.org/IMGTrepertoire/index.php?section=LocusGenes&repertoire=genetable&species=human&group=TRAV). Only MHCI predictions are available, as mentioned in the webserver. Even though there is no MHC input, the maximum peptide length allowed is 12 (as per the webserver instructions), which is smaller than all MHCII peptide lengths with associated TCR sequence in our benchmarking dataset. The prediction also needs CDR3$\alpha\beta$ sequences. We submitted the TCR and peptide sequences to the webserver at https://services.healthtech.dtu.dk/services/NetTCR-2.2/ [63] with the default parameters Similarity scaling factor: 10 and Percentile rank threshold: 100.

#### ImRex

We downloaded the source code and pre-trained models from https://github.com/pmoris/ImRex/tree/master [38] and ran them locally, using the same pretrained model and Python command line as specified in the 'Predictions using the pre-built model' section of the GitHub README file. Prediction was available only for TCRs with CDR3$\beta$ sequences. We used the output TCR-pMHC prediction score to calculate the AUCs in Figure 2b. While, running the above, we received the following output messages in terminal: "Filtered CDR3 sequences to length: (10, 20)" and "Filtered epitope sequences to length: (8, 11)", indicating prediction restrictions.

#### ERGO II

We downloaded the source code from the main GitHub link https://github.com/IdoSpringer/ERGO-II/tree/master [41, 64] and ran it locally using the same Python command line as specified in the 'Prediction Instructions' section of the GitHub README file. Note that the files Model.py and Predict.py required some changes before they could be run, including renaming a directory "TCR_Autoencoder" in the code to "Models/AE" since the TCR_autoencoder directory was missing. We also added the specification *cpu-only* in load-torch settings. On our benchmarking dataset, the ERGO model trained on the McPAS database performed better than the model trained on the VDJdb database, so we reported the former model scores. Prediction was available only for TCRs with CDR3$\beta$ sequences.

#### epiTCR

We downloaded the source code from the GitHub link: https://github.com/ddiem-ri-4D/epiTCR [40] and ran it locally using the same pretrained model and Python command line as specified in the 'Run prediction using the pre-trained model' section of the GitHub README file. Prediction was available only for TCRs with CDR3$\beta$ sequences. The code requires MHC allele sequences, which were copied from the sample test file entries ("MHC" column) corresponding to the same alleles, downloaded from https://github.com/ddiem-ri-4D/epiTCR/blob/main/data/hlaCovertPeudoSeq/HLAWithPseudoSeq.csv.zip. We used the output TCR-pMHC predicted binding probability ('predict_proba' column in the output file).

While running the epiTCR package locally, we received the following warning: "UserWarning: Trying to unpickle estimator DecisionTreeClassifier from version 1.2.0 when using version 1.1.2. This might lead to breaking code or invalid results. Use at your own risk. For more info please refer to: https://scikit-learn.org/stable/model_persistence.html#security-maintainability-limitations." Furthermore, [40] mentions: "CDR3$\beta$ sequences of 8–19 amino acids were provided as TCR input, and peptide sequences of 8–11 amino acids were given as the corresponding peptide." The code returns as error if the lengths exceed this limit, so MHCII and TIL1383I TCR predictions could not be done.

#### iTCep

We downloaded the source code from the GitHub link: https://github.com/kbvstmd/iTCep/ [65] and ran it locally. For peptides without a corresponding CDR3$\beta$ sequence, the model predicts a list of possible binding TCRs sequences. We do not use these predictions. [65] mentions "...the peptide-CDR3 pair sequences were padded to the maximum length of 32 and..." Longer pairs in the input data did not return any result in the output file, so no MHCII predictions were available.

#### Attn-TAP

We downloaded the source code from the GitHub link: https://github.com/Bioinformatics7181/AttnTAP [66] and ran it locally. Prediction was available only for TCRs with CDR3$\beta$ sequences. The mcpas model performed better than the vdjdb model on our benchmarking dataset, so we reported the former model scores.

#### ATM-TCR

We downloaded the source code from the GitHub link: https://github.com/Lee-CBG/ATM-TCR [51] and ran it locally. Prediction was available only for TCRs with CDR3$\beta$ sequences.

#### DeepTR

Only HLAI predictions were available, of them, no HLA-B81:01 prediction was available. We uploaded peptide data for individual TCRs separately to the webserver at https://bioinfo.uth.edu/DeepTR/Prediction.php [67] and downloaded the results. Prediction was available only for TCRs with full sequences available.

#### TCRPrediction

We downloaded the source code from the GitHub link: https://github.com/kyoheikoyama/TCRPrediction [68] and ran it locally. Prediction was available only for TCRs with CDR3$\beta$ sequences.

#### HeteroTCR

We downloaded the source code from the GitHub link: https://github.com/yuzilan/HeteroTCR [69] and ran it locally. Prediction was available only for TCRs with CDR3$\beta$ sequences and for peptides with length <15.

#### TITAN

We downloaded the source code from the GitHub link: https://github.com/PaccMann/TITAN [45] and ran it locally. Prediction was available only for TCRs with full sequences available.
